# Targeted cancer therapy induces APOBEC fuelling the evolution of drug resistance

**DOI:** 10.1101/2020.12.18.423280

**Authors:** Manasi K. Mayekar, Deborah R. Caswell, Natalie I. Vokes, Emily K. Law, Wei Wu, William Hill, Eva Gronroos, Andrew Rowan, Maise Al Bakir, Caroline E. McCoach, Collin M. Blakely, Nuri Alpay Temiz, Ai Nagano, D. Lucas Kerr, Julia K. Rotow, Franziska Haderk, Michelle Dietzen, Carlos Martinez Ruiz, Bruna Almeida, Lauren Cech, Beatrice Gini, Joanna Przewrocka, Chris Moore, Miguel Murillo, Bjorn Bakker, Brandon Rule, Cameron Durfee, Shigeki Nanjo, Lisa Tan, Lindsay K. Larson, Prokopios P. Argyris, William L. Brown, Johnny Yu, Carlos Gomez, Philippe Gui, Rachel I. Vogel, Elizabeth A. Yu, Nicholas J. Thomas, Subramanian Venkatesan, Sebastijan Hobor, Su Kit Chew, Nnennaya Kanu, Nicholas McGranahan, Eliezer M. Van Allen, Julian Downward, Reuben S. Harris, Trever G. Bivona, Charles Swanton

## Abstract

The clinical success of targeted cancer therapy is limited by drug resistance that renders cancers lethal in patients^1-4^. Human tumours can evolve therapy resistance by acquiring *de novo* genetic alterations and increased heterogeneity via mechanisms that remain incompletely understood^1^. Here, through parallel analysis of human clinical samples, tumour xenograft and cell line models and murine model systems, we uncover an unanticipated mechanism of therapy-induced adaptation that fuels the evolution of drug resistance. Targeted therapy directed against EGFR and ALK oncoproteins in lung cancer induced adaptations favoring apolipoprotein B mRNA-editing enzyme, catalytic polypeptide (APOBEC)-mediated genome mutagenesis. In human oncogenic *EGFR*-driven and *ALK*-driven lung cancers and preclinical models, EGFR or ALK inhibitor treatment induced the expression and DNA mutagenic activity of *APOBEC3B* via therapy-mediated activation of NF-κB signaling. Moreover, targeted therapy also mediated downregulation of certain DNA repair enzymes such as UNG2, which normally counteracts APOBEC-catalyzed DNA deamination events. In mutant *EGFR*-driven lung cancer mouse models, APOBEC3B was detrimental to tumour initiation and yet advantageous to tumour progression during EGFR targeted therapy, consistent with TRACERx data demonstrating subclonal enrichment of APOBEC-mediated mutagenesis. This study reveals how cancers adapt and drive genetic diversity in response to targeted therapy and identifies APOBEC deaminases as future targets for eliciting more durable clinical benefit to targeted cancer therapy.

## Main Text

Cancer initiation, evolution and progression is controlled, in part, by the acquisition of genetic alterations through a variety of mechanisms. Molecularly targeted therapy against specific driver alterations such as mutant EGFR and ALK gene rearrangements in lung cancer has dramatically improved outcomes for patients^2-5^. However, the development of targeted therapy resistance remains an unresolved challenge and a barrier to maximizing clinical success^6^. Resistance can occur due to the outgrowth of preexisting resistant clones or due to *de novo* acquisition of new mutations in cancer cells^1^. Tumour genetic diversity and mutational burden is elevated in drug resistant EGFR-mutant lung adenocarcinomas, one of the major molecular subtypes of lung adenocarcinomas^7^. Higher tumour mutational burden correlates with poor clinical outcome in such patients. Moreover, genome instability mechanisms contributing to the evolution of resistance remain unclear. Hence, new approaches that can block the emergence of genetic diversity and resistance are urgently needed.

The APOBEC family of enzymes converts single-stranded DNA cytosines to uracils. Unrepaired uracil lesions can template the insertion of adenine nucleobases and become immortalized through replication as C-to-T mutations. Humans have seven APOBEC3 family members (APOBEC3A-H) and all but one (G) preferentially deaminates TC dinucleotide substrates^8^. Prior tumour genome sequencing work including TRACERx revealed an enrichment for mutations in an APOBEC preferred substrate context subclonally and later in non-small cell lung cancer (NSCLC) evolution^9-12^. Additionally, certain targeted therapy-resistance conferring mutations involve C-to-T mutations and a subset of those occur in the APOBEC-preferred TC context^8,12,13^.

Based on these collective observations, we hypothesized that APOBEC-mediated genome mutagenesis could facilitate genetic adaptation in response to targeted therapy and contribute to the acquisition of genetic diversity and drug resistance. We tested this hypothesis in paradigm-defining models of targeted cancer therapy: oncogenic *EGFR*-driven and *EML4-ALK*-driven models of lung adenocarcinoma treated with EGFR or ALK tyrosine kinase inhibitors (TKIs).

### Targeted therapy induces adaptations favoring APOBEC-mediated genome mutagenesis

To test the hypothesis, we first examined targeted therapy-induced transcriptional changes in human *EGFR*-mutant lung adenocarcinoma systems (patient-derived cellular models) present in publicly available datasets (GEO2R). We noted that treatment with the EGFR inhibitor erlotinib was associated with transcriptional upregulation of certain *APOBEC3* subfamily genes and downregulation of the repair factor *UNG*, the key enzyme required for repair of APOBEC-induced uracil lesions, both acutely and at later time points (Extended Data Fig. 1a).

To confirm these transcriptional changes, we performed RNA-Seq analysis on cells generated from an established patient-derived cell line model of oncogenic *EGFR*-driven lung adenocarcinoma, PC9 cells (*EGFR exon19del*), that were exposed to EGFR TKI treatment for a sustained timeframe. We found that these EGFR TKI-treated cells showed significant upregulation of *APOBEC3B, APOBEC3C* and *APOBEC3F* transcripts and downregulation of *UNG* transcripts (Extended Data Fig. 1b). We noted that the transcriptional effects identified were a conserved consequence of EGFR pharmacologic inhibition in *EGFR*-driven lung adenocarcinomas and not specific to the class of EGFR inhibitor (Extended Data Fig. 1a). Transcript levels of other *APOBEC3* family members, including *activation-induced cytosine deaminase* (*AID*), were minimally detected (TPM<1.5). We validated these findings in both oncogenic *EGFR*-driven and *EML4-ALK*-driven cellular models of lung adenocarcinoma at both the RNA and protein level (Fig. 1a-c and Extended Data Fig. 1c-e). We next validated these results at the functional level using established assays to quantify APOBEC activity^14^ and uracil excision capacity^15^. Consistent with RNA and protein level validations, TKI treatment resulted in a marked increase in nuclear APOBEC activity (Fig. 1d, Extended Figure 1f-h), and a decrease in nuclear uracil excision capacity in multiple *EGFR*-driven and *EML4-ALK*-driven models of lung adenocarcinoma (Fig. 1e, Extended Data Figure 1i-k).

**Fig. 1.**
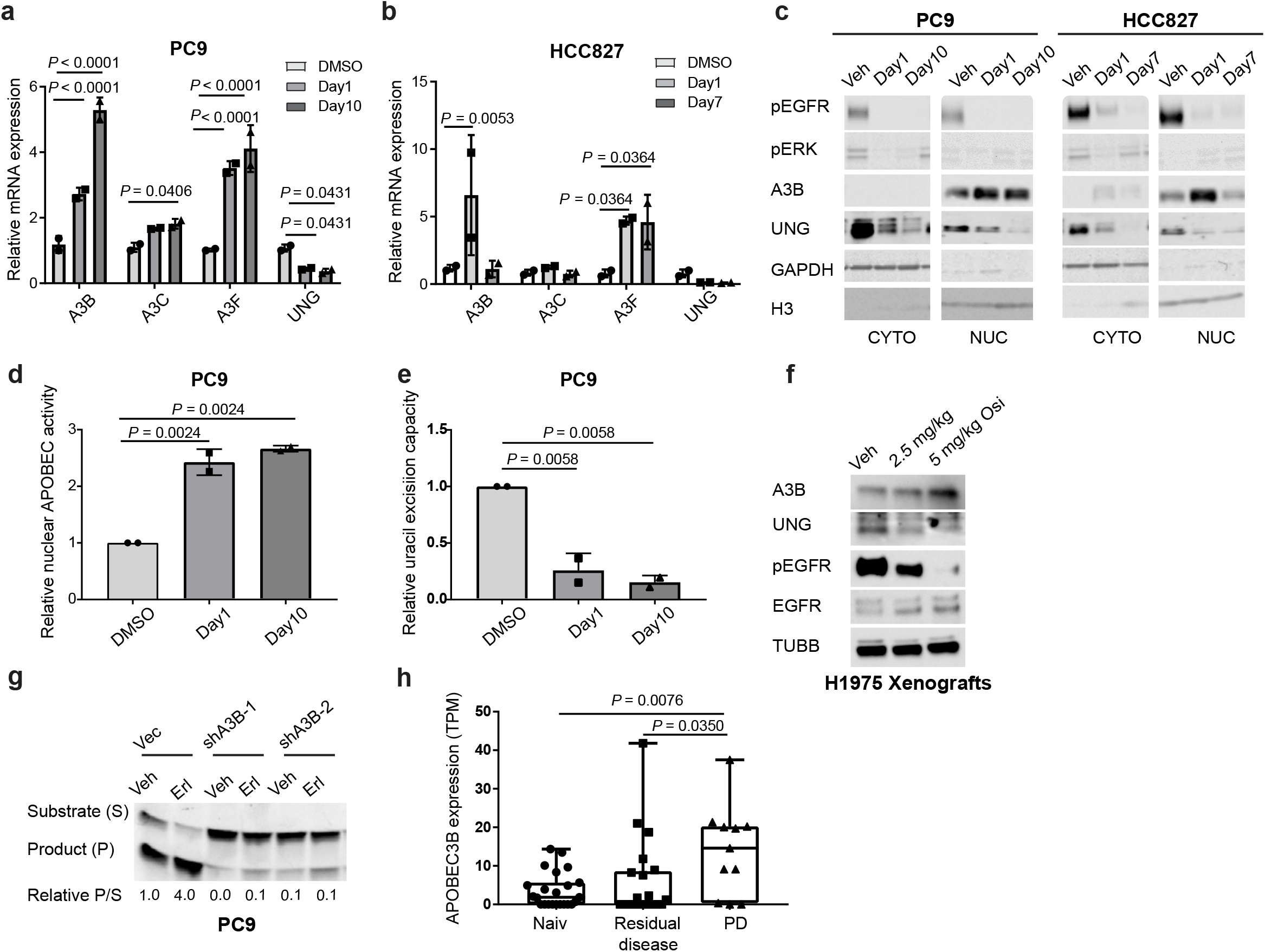
Treatment with EGFR inhibitors induces adaptations favorable for APOBEC-mediated mutagenesis. **a, b**, RT-qPCR analysis of RNA from PC9 cells were treated with DMSO or 2 μM osimertinib (osi) for 18 hours or 2 μM osimertinib for 10 days. HCC827 cells were treated with DMSO or 0.4 μM osimertinib for 18 hours or 0.4 μM osimertinib for 7 days (n = 2 biological replicates, mean ± SD, two-way ANOVA test). **c**, Western blot analysis of cells treated in a similar manner as in **a** and **b** (CYTO: cytoplasmic extracts, NUC: nuclear extracts). **d, e**, APOBEC activity assay (**d**) and uracil excision capacity (**e**) assay performed using nuclear extracts of PC9 cells treated similarly as indicated for panel **a** (n = 2 biological replicates, mean ± SD, ANOVA test). **f**, Western blot analysis using extracts of *EGFR*-mutant H1975 human NSCLC xenografts harvested after 4 days of treatment with vehicle or the indicated doses of osimertinib. **g**, APOBEC activity assay performed using nuclear extracts of PC9 cells transduced with pLKO vector or its derivatives encoding shRNAs against APOBEC3B and treated with DMSO or 1 μM erlotinib (erl) for 18 hours. **h**, Comparison of APOBEC3B expression levels measured using RNA-Seq analysis in human NSCLC specimens obtained before treatment (naive), or on-treatment at residual disease (RD) or at progressive disease (PD) stages from lung cancer patients undergoing treatment with tyrosine kinase inhibitors (whiskers: min to max, ANOVA test). (H3: Histone H3, TUBB: Tubulin Beta Class I).

Because the expression of APOBEC and DNA repair enzymes can be coupled to the cell cycle, which is suppressed by TKI therapy, we treated cells with a CDK4/6 cell cycle inhibitor palbociclib^16^ and measured *APOBEC* and *UNG* expression. While *UNG2* expression decreased upon palbociclib treatment, we also detected a substantial decline in *APOBEC3B* expression (Extended Data Fig. 1l). This suggests that TKI-mediated induction of APOBEC is unlikely to be a consequence of TKI treatment-induced cell cycle inhibition.

We next examined whether these findings extended to *in vivo* models of human lung adenocarcinoma. An increase in APOBEC3B protein levels and a decrease in UNG protein levels were detected in tumour tissues obtained from oncogenic *EGFR*-driven tumour xenograft models of human lung adenocarcinoma treated with the EGFR TKI osimertinib (Fig. 1f and Extended Data Fig. 1m). Additionally, RNA-seq results from a patient-derived tumour xenograft (PDX) model of lung adenocarcinoma (EGFR L858R)^17^ treated with erlotinib revealed an increase in *APOBEC3B* mRNA levels and a decrease in *UNG2* mRNA levels upon erlotinib treatment (Extended Data Fig. 1n). The collective findings support one plausible model whereby EGFR or ALK-targeted therapy induces adaptive conditions in cancer cells that may be favorable for APOBEC3-mediated genome mutagenesis, a model we further investigated.

### APOBEC3B is a driver of TKI-induced nuclear APOBEC activity

We next investigated which APOBEC3 family member drives an increase in nuclear APOBEC activity in response to TKI treatment. Osimertinib treatment resulted in an increase in nuclear APOBEC activity both acutely and at later stages of continuous treatment (Extended Data Fig. 2a-b). We examined which APOBEC family member is upregulated upon TKI treatment during these time points (Fig. 1a, Extended Data Fig. 1b). Only the expression of APOBEC3B closely mirrored the changes in nuclear APOBEC activity that we observed (Extended Data Fig. 2c-f). Accordingly, depletion of APOBEC3B in PC9 (EGFR exon19del) or H3122 cells (EML4-ALK) using RNAi or CRISPR-Cas9 mediated approaches, respectively, resulted in a substantial reduction in both baseline and TKI-induced nuclear APOBEC activity (Fig. 1g, Extended Data Fig. 2g-i). These results implicate APOBEC3B as a key driver of TKI-induced nuclear APOBEC activity (in addition to baseline activity)

Next, we assessed the clinical relevance of our initial findings by examining APOBEC3B expression in clinical specimens of non-small cell lung cancer (NSCLC) obtained from patients before (treatment naïve, TN) or during targeted therapy, at residual disease (RD) during an initial treatment response, or at progressive disease (PD) during continuous treatment (Supplementary Table 1)^18^. Human tumours that were exposed to targeted therapy showed increased *APOBEC3B* mRNA expression, particularly at PD (Fig. 1h). Thus, APOBEC3B expression was elevated both in patient-derived models of oncogene-driven lung cancer and in human tumours exposed to targeted therapy.

### APOBEC3B is detrimental at tumour initiation

Mouse models are a key system for the advancement of our understanding of lung cancer targeted therapy resistance^11,19-21^. However, recent findings demonstrate that these models lack the mutational heterogeneity present in human tumours^22-24^. This may be due, in part, to the fact that mice encode only a single, cytoplasmic and non-genotoxic APOBEC3 enzyme^25,26^. To address this, we combined a new Cre-inducible model for human *APOBEC3B* expression (*Rosa26::LSL-A3Bi*) with an established EGFR^L858R^ driven lung cancer mouse model^19,20,27^, and a Cre-inducible tetracycline controlled transactivator (*R26*^*LNL-tTA*^). We induced tumours in *TetO-EGFR*^*L858R*^*;R26*^*LNL-tTA/+*^ *(E)* or *TetO-EGFR*^*L858R*^*;R26*^*LNL-tTA/LSL-APOBEC3B*^ (*EA3B)* mice (Fig 2a), and at three months after induction found that the proportion of mice with tumours, and total tumour volume per mouse was significantly higher in *E* than *EA3B* mice (Fig 2b and c).

**Fig. 2.**
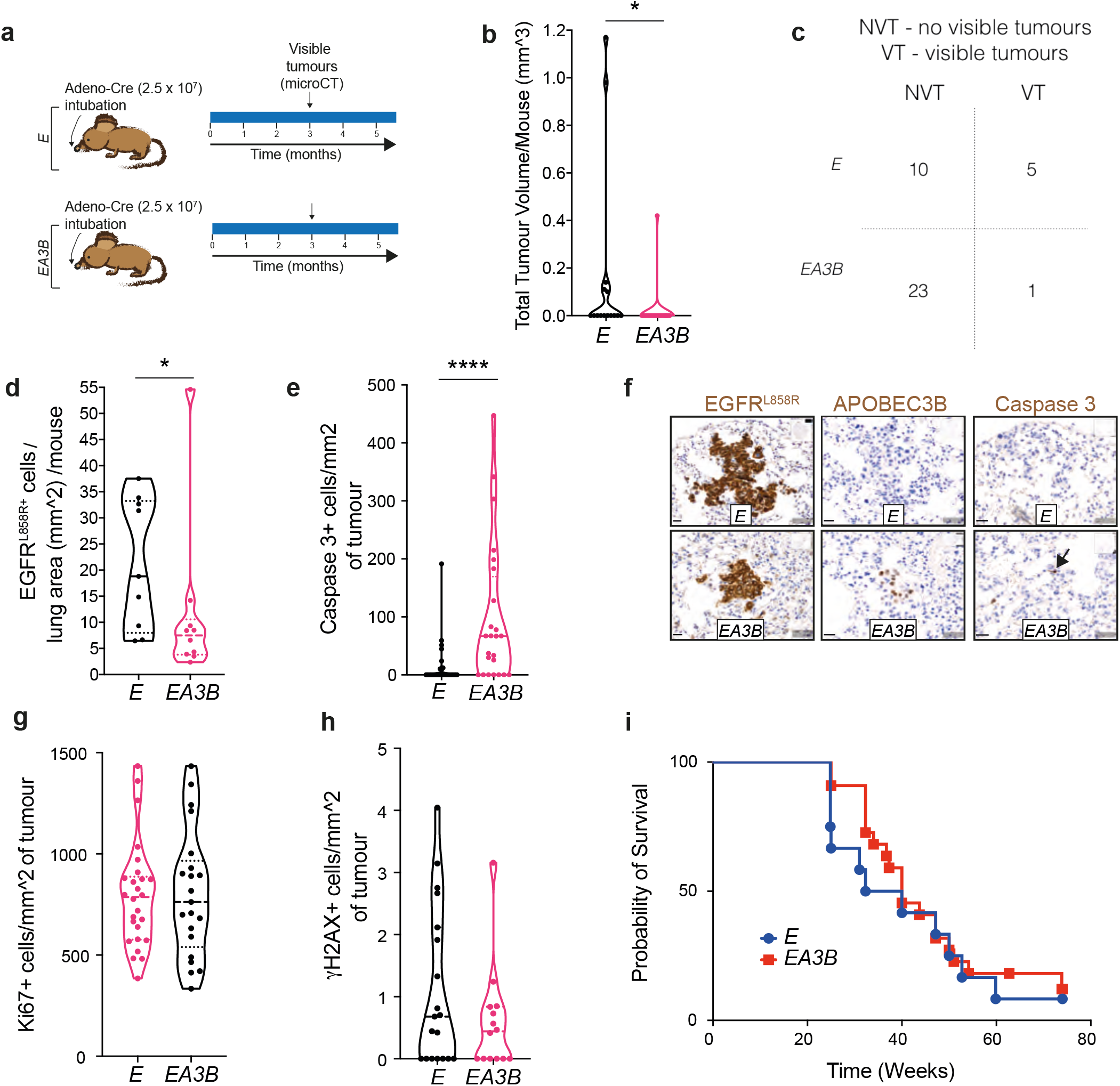
APOBEC3B is detrimental for tumourigenesis in a mouse model of EGFR^L858R^ driven lung cancer. **a**, Tumours in *TetO-EGFR*^*L858R*^*;R26*^*LNL-tTA/+*^ (*E*) and *TetO-EGFR*^*L858R*^*;R26*^*LNL-tTA/LSL-APOBEC3B*^ (*EA3B*) mice were induced using the indicated viral titer (Adeno-Cre). Tumour growth was assessed by microCT. **b**, Total tumour volume per mouse at 3 months post induction quantified by microCT analysis (*E* = 15, *EA3B* = 24, median = dashed line, 1st and 3rd quartiles = dotted lines, two-sided Mann-Whitney Test, P = 0.0163, each dot represents a mouse). **c**, 2×2 Contingency table of the number of mice with visible or no visible tumours by microCT at 3 months post induction (two-sided Fisher’s exact test, P<0.05). **d**, Quantification of EGFR^L858R^ positive cells per lung area by immunohistochemical (IHC) staining at 3 months post induction (*E* = 9, *EA3B* = 10, median = dashed line, 1st and 3rd quartiles = dotted lines, two-sided Mann-Whitney Test, P =0.0435, each dot represents a mouse). **e**, Quantification of Caspase 3 + cells per mm^2 of tumour at 3 months post induction (*E* = 9, *EA3B* = 10, median = dashed line, 1st and 3rd quartiles = dotted lines, two-sided Mann-Whitney Test, P<0.0001, each dot represents a tumour) **f**, Representative IHC staining of EGFR^L858R^, APOBEC3B, and Caspase 3. Scale bar = 20 um, arrow indicates positive cells. **g**, Quantification of Ki67 positive cells per mm^2 of tumour at 3 months post induction (*E* = 9, *EA3B* = 10, each dot represents a tumour). **h**, Quantification of γH2AX positive cells per mm^2 of tumour at 3 months post induction (*E* = 9, *EA3B* = 10, each dot represents a tumour) **i**, Survival curve of *E* versus *EA3B* mice (*E* =15, *EA3B* = 24, each dot represents a tumour).

To verify this result, an additional set of mice was induced and then culled at 3 months. The lungs were removed and fixed for immunohistochemical (IHC) staining. EGFR^L858R^ staining revealed a significantly increased number of EGFR^L858R^ positive cells per lung area per mouse in *E* control mice compared to *EA3B* experimental mice, substantiating the likely detrimental effect of APOBEC3B early in tumourigenesis (Fig 2d).

Staining for the molecular marker of DNA damage gH2AX and the proliferation marker Ki67 revealed no significant differences between *E* and *EA3B* tumours, while staining for the programmed cell death marker caspase 3 showed a significant increase in *EA3B* mouse tumour cells, suggesting that *APOBEC3B* expression in tumours at this early stage leads to increased tumour cell death (Fig 2e-h). Overall survival of the animals was not significantly different in *E* vs *EA3B* mice (Fig 2i).

### APOBEC3B drives TKI resistance in an EGFR^L858R^ lung cancer mouse model

Based on our findings that APOBEC3B was detrimental for tumour initiation and findings from the TRACERx 100 dataset that APOBEC-mediated mutagenesis is enriched subclonally, later in tumour evolution in *EGFR*-mutant disease (Extended Data Fig 3a-b) and the wider cohort^10^, we generated mice in which *APOBEC3B* expression could be temporally separated from *EGFR*^*L858R*^. This allowed us to mimic the subclonal acquisition of APOBEC mutations in NSCLC reflective of the role of APOBEC in human NSCLC. We used a tetracycline-inducible pneumocyte specific mouse model of EGFR-dependent lung cancer^19,20,27^ combined with a Cre recombinase-steroid receptor fusion (Rosa26CreER^T2^)^28^, and the conditional human *APOBEC3B* minigene used above to generate *CCSP-rtTA;TetO-EGFR*^*L858R*^*;Rosa26*^*CreER(T2)/LSL-APOBEC3B*^ (*EA3Bi*) mice (Fig 3a). *EGFR*^*L858R*^ was induced in mice with doxycycline containing chow, and after 6 weeks mice were placed on cyclical TKI treatment regimen (erlotinib, 25 mg/kg, 5 days per week) previously shown to induce rapid resistance in this model^20^ (Fig. 3a). One week after initiation of TKI therapy, *APOBEC3B* expression was induced by administration of tamoxifen (150 mg/kg, 3 times over 1 week) (Fig. 3a). After 2 cycles of TKI treatment with one treatment holiday (Figure 3a) at 5 months after tumour induction, all mice were culled and the lungs were harvested.

**Fig. 3.**
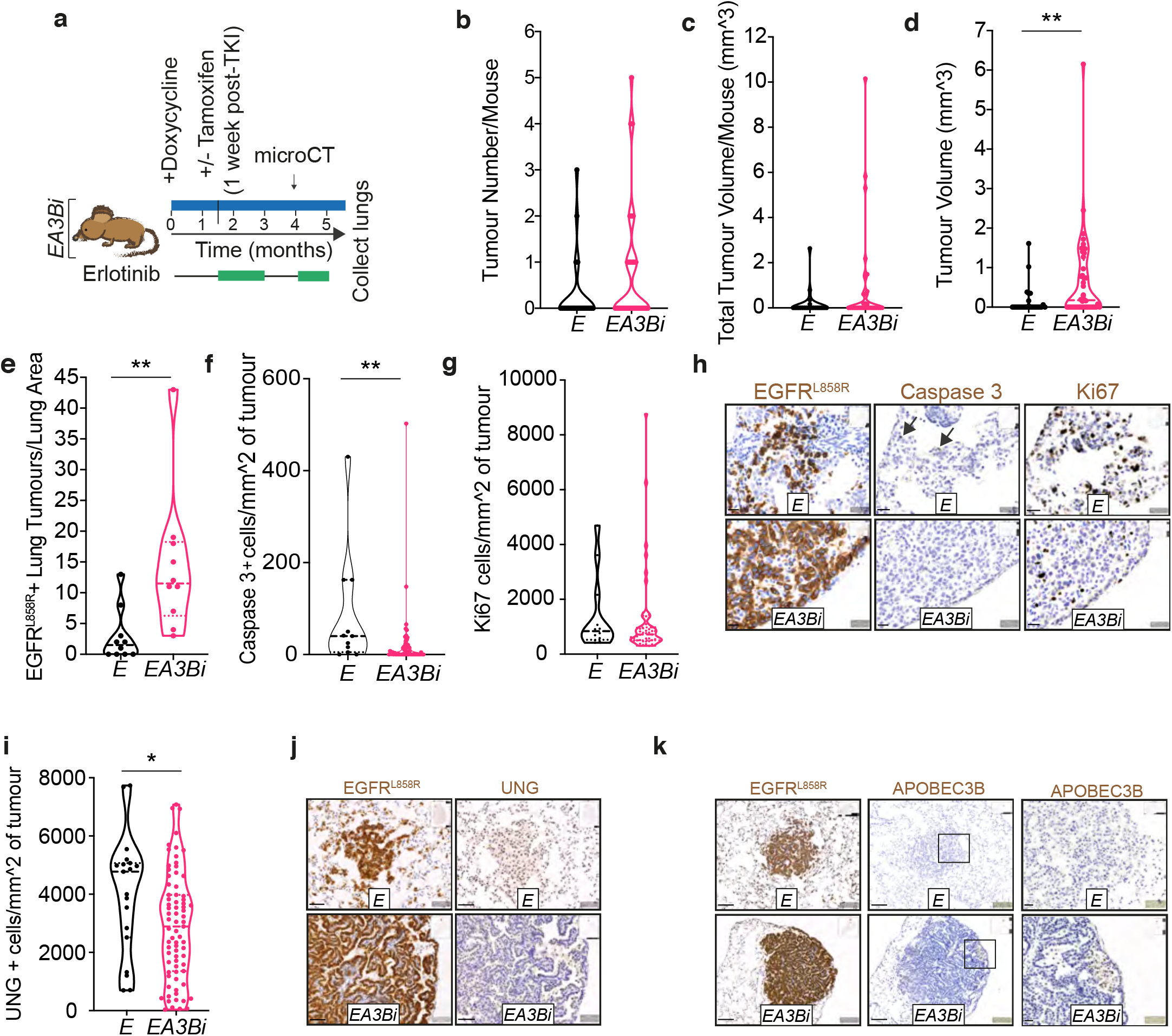
APOBEC3B drives TKI resistance in an EGFR^L858R^ mouse model of lung adenocarcinoma. **a**, Experimental set up of induction of APOBEC3B mutagenesis post-TKI using *TetO-EGFR*^*L858R*^*;CCSP-rt-TA;R26*^*LSL-APOBEC3B/Cre-ER(T2)*^*(EA3Bi)* mice. **b**, Tumour number per mouse at 4 months post induction (*E* n = 23, *EA3Bi* n = 31, each dot represents a mouse). **c**, Total tumour volume (mm^3) per mouse at 4 months post induction (*E* n = 23, *EA3Bi* n = 31, each dot represents a mouse) **d**, Tumour volume (mm^3) at 4 months post induction (*E* n = 23, *EA3Bi* n = 31, median = dashed line, 1st and 3rd quartiles = dotted lines, two-sided Mann-Whitney Test, P = 0.0075, each dot represents a tumour). **e**, Quantification of EGFR^L858R+^ lung tumours per lung area at 5 months post induction (*E* n=10, *EA3Bi* n=10, median = dashed line, 1st and 3rd quartiles = dotted lines, two-sided Mann-Whitney Test, P = 0.0012, each dot represents a mouse). **f**, Quantification of Caspase 3+ cells per mm^2 of tumour at 5 months post induction (*E* n=6, *EA3Bi* n =6, median = dashed line, 1st and 3rd quartiles = dotted lines, two-sided Mann-Whitney Test, P = 0.0082, each dot represents a tumour). **g**, Ki67+ cells per mm^2 of tumour at 5 months post induction (*E* n=10, *EA3Bi* n=10, each dot represents a tumour) **h**, Representative serial sections of EGFR^L858R^, Caspase 3, Ki67. Scale bar = 20 um, arrows indicate positive cells. **i**, Quantification of UNG+ cells per mm^2 of tumour at 5 months post induction (*E* n=10, *EA3Bi* n=10, median = dashed line, 1st and 3rd quartiles = dotted lines, two-sided t-test, P = 0.0226, each dot represents a tumour). **j**, Representative immunohistochemical staining of EGFR^L858R^ and UNG. Scale bar = 50 um **k**, Representative immunohistochemical staining of EGFR^L858R^ and APOBEC3B. Scale bar = 100 um and 20 um.

MicroCT analysis at 4 months after tumour induction revealed a higher tumour number and total tumour volume per mouse, and a significantly higher average tumour volume in *EA3Bi* mice compared to *E* mice (Fig 3b-d). Immunohistochemical (IHC) staining of the lungs for *EGFR*^*L858R*^ at 5 months post tumour induction after the second treatment cycle revealed a significantly higher number of tumours per lung area in *EA3Bi* mice compared to *E* mice (Fig 3e).

IHC staining for caspase 3 demonstrated that *EA3Bi* mice had significantly lower levels of cancer cell death per tumour (Fig 3f). There was no difference in Ki67 staining between *E* an *EA3Bi* cohorts (Fig 3g-h). These data suggest that tumours in *E* mice are more sensitive to TKI therapy than *EA3Bi* tumours, suggesting that APOBEC3B contributes to increased TKI resistance. Serial sections stained for EGFR^L858R^ and APOBEC3B revealed heterogeneous APOBEC3B expression in *EA3Bi* tumours (Fig 3k). UNG staining revealed a significant decrease in UNG positive cells per tumour in *EA3Bi* mice treated with TKI therapy, aligning with our human preclinical findings (Fig 1, and Extended Fig 1, and Fig 3i and j).

### NF-κB signaling is critical for TKI-induced APOBEC3B upregulation

We next investigated the mechanism by which TKI treatment induces APOBEC3B upregulation using the human lung cancer systems. Our previous study revealed that NF-κB signaling is activated upon EGFR oncogene inhibition in human lung cancer, perhaps as a stress response^17^. Other studies suggest that NF-κB signaling may be a prominent driver of *APOBEC3B* gene expression^29,30^. To test whether NF-κB signaling promotes *APOBEC3B* expression during targeted therapy, we examined our RNA-seq dataset generated from *EGFR*-driven human lung adenocarcinoma cells treated acutely with erlotinib and an established NF-κB inhibitor alone or in combination^17^. We found that TKI treatment induced transcriptional upregulation of *APOBEC3B* could be attenuated by co-treatment with an established NF-κB inhibitor^17^ (Extended Data Fig. 4a). We next examined the changes in the nuclear NF-κB (RELA and RELB subunits) levels upon TKI treatment and found that an increase in nuclear RELB tracked more consistently with the increase in APOBEC3B levels (Extended Data Fig. 4b). These data suggest that the NF-κB pathway may be a driver of *APOBEC3B* expression. To confirm this, we induced NF-κB pathway with increasing doses of TNFα, which increased nuclear RELA and RELB levels as expected and induced APOBEC3B as well (Extended Data Fig. 4c-d). Furthermore, inhibition of the NF-κB pathway by depletion of RELA or RELB reduced baseline and TKI-induced APOBEC3B levels (Extended Data Fig. 4e-f). These data uncover NF-κB activation resulting from EGFR TKI treatment as a driver of APOBEC3B upregulation in response to therapy and suggest functional cooperativity or redundancy between RELA and RELB in promoting TKI-induced APOBEC3B expression.

To determine the clinical relevance of these findings, we examined single cell RNA Seq data obtained from cancer cells in tumours obtained from patients with NSCLC before or while on targeted therapy^18^. We observed that, like APOBEC3B, the mRNA expression of both RELA and RELB was significantly increased in tumours exposed to EGFR TKI treatment, particularly at tumour progression (Fig. 1h and Extended Data Fig. 4g-i).

### *UNG2* downregulation is associated with c-Jun suppression during TKI treatment

We investigated the potential mechanism by which *UNG2* mRNA expression is transcriptionally downregulated during TKI treatment. *UNG* gene promoter analysis (using PROMO)^31^ revealed the presence of predicted c-Jun consensus binding sites. Our RNA Seq data from EGFR TKI-treated PC9 EGFR mutant NSCLC cells indicated that like *UNG, c-JUN* was also transcriptionally downregulated upon TKI treatment, which we validated using RT-qPCR analysis (Extended Data Fig. 4j-k). These data are consistent with the expectation that c-Jun expression should decrease upon inhibition of the MAPK pathway that occurs as a result of EGFR inhibition by TKI treatment^32^. Thus, we investigated whether *UNG* downregulation that occurs upon TKI treatment might be caused by *c-JUN* downregulation that results from TKI treatment. Consistent with this hypothesis, c-Jun depletion alone was sufficient to suppress *UNG2* expression, suggesting that *UNG* downregulation may be a consequence, in part, of the *c-JUN* suppression that occurs as a consequence of TKI treatment (Extended Data Fig. 4k).

### APOBEC mutational signature is acquired during targeted therapy in an APOBEC3B-dependent manner in clonal cellular models of human lung cancer

Our findings suggested that *APOBEC3B* upregulation (coupled with *UNG2* downregulation) could promote genome mutagenesis during therapy. We tested whether APOBEC signature cancer genome mutations are acquired during TKI treatment in oncogenic *EGFR*- and *EML4-ALK*-driven lung adenocarcinoma models. Analysis of whole exome and whole genome sequencing (WES and WGS) data from our cell line models and from prior studies^33,34^ revealed that TKI treatment was associated with the acquisition of APOBEC signature mutations in multiple models during TKI treatment and in TKI-resistant derivatives of PC9 cells including those with established resistance mutations in *PIK3CA, BRAF* and *NRAS* genes in the APOBEC-preferred nucleotide context and those with EGFR T790M resistance mutation^35^(Extended Data Fig. 5a-e).

To evaluate the impact of APOBEC3B on response to targeted therapy, we depleted APOBEC3B in both clonal and polyclonal populations using RNAi or CRISPR-Cas9 technologies and observed that APOBEC3B-proficiency was beneficial to lung adenocarcinoma cells during long-term treatment with TKIs in both oncogenic *EGFR*-driven and *EML4-ALK*-driven cellular models (Extended Data Fig. 6).

To further investigate the effect of APOBEC3B on tumour evolution during treatment, we treated APOBEC3B-proficient and APOBEC3B-deficient isogenic clonal models of PC9 cells for up to 2.5 months with osimertinib. A nuclear APOBEC activity assay confirmed the absence of both baseline and TKI-induced nuclear APOBEC activity in APOBEC3B-deficient cells (Extended Data Fig. 7a). We found that the APOBEC3B-proficient cells showed acquisition of advantageous signaling alterations such as higher STAT3 and AKT activation after long-term osimertinib treatment (Extended Data Fig. 7b). Furthermore, our mutational signature analysis revealed that TKI treatment drives increased genetic heterogeneity and acquisition of an APOBEC mutational signature in APOBEC3B-proficient but not APOBEC3B-deficient cells (Extended Data Fig. 7c-d). These data support the conclusion that APOBEC3B drives tumour evolution through genetic adaptation during TKI treatment that is advantageous for tumour cell survival and the development of resistance.

### APOBEC mutagenesis is upregulated during targeted therapy in human lung cancer clinical specimens along with the acquisition of TKI resistance-conferring mutations

To extend the clinical relevance of our preclinical findings, we performed mutational signature analysis on WES data obtained from tumours collected from treatment naïve and TKI-treated lung cancer patients (Supplementary Table 2). We focused on mutational signatures most commonly observed in lung cancer patients including signatures 1, 4 and 5 that are predicted to be induced by deamination of 5-methylcytosines, smoking and an unknown mechanism respectively; while signatures 2 and 13 are predicted to be induced due to APOBEC activity^36^. This analysis revealed that a high proportion of APOBEC-associated mutations were more common in a post-TKI setting and were substantially enriched after TKI treatment in certain patients (Fig. 4a-b). Overall, the proportion of APOBEC mutational signatures detected in the tumours showed an increasing trend in tumours that were exposed to targeted therapy (Extended Data Fig. 8). We also identified APOBEC signature mutations that could contribute to resistance by AKT pathway activation such as an activating mutation in *PIK3CA* (E545K)^37^ and inactivating mutation in *PTEN* (S287*) or MAPK pathway reactivation by inactivation of *PP2A*, a negative regulator of MAPK signaling^38^, and an ALK inhibitor desensitizing mutation in *ALK* (E1210K)^39^ in the tumours of some patients who had progressed on or shown incomplete response to EGFR inhibitor therapy (Supplementary Tables 2-3). AKT and MAPK pathway activation are known to cause EGFR and ALK inhibitor resistance^38,40-44^.

**Fig. 4.**
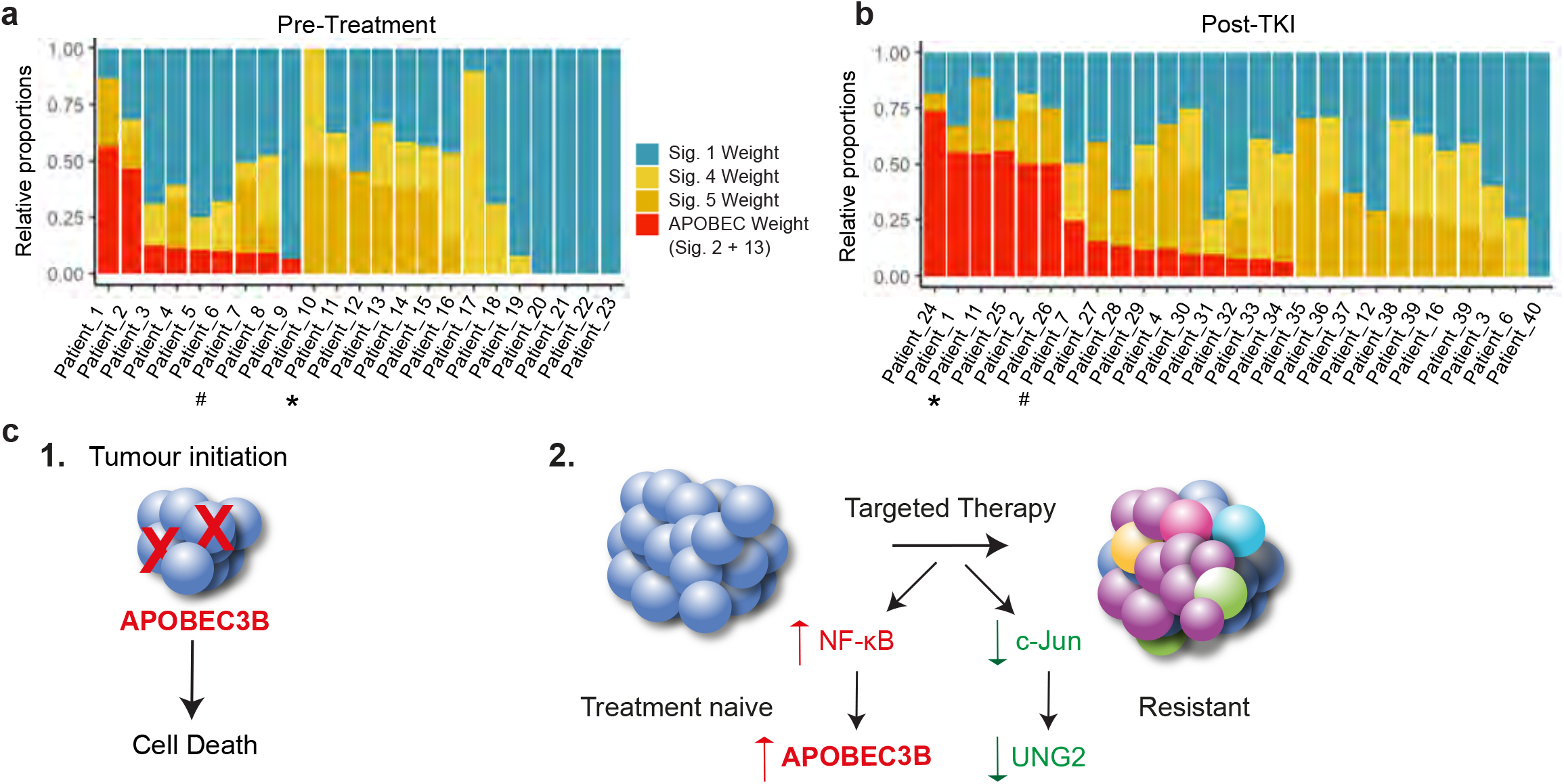
Mutational signature analyses of *EGFR*- and *ALK*-mutant NSCLC clinical specimens pre- and post-TKI treatment. **a, b**, Analyses showing the proportion of mutational signatures in whole exome sequencing data obtained from *EGFR* and *ALK*-mutant NSCLC patients (**a**) pre- and (**b**) post-TKI treatment. *, #: Indicate patients whose tumours showed substantial increase in the proportion of APOBEC-mediated mutations after TKI treatment compared to their tumor biopsies taken prior to TKI treatment. **c**, Model for the role of APOBEC3B in tumour initiation and targeted cancer therapy-induced adaption during the evolution of drug resistance. APOBEC3B is detrimental at the tumour initiation stage but fuels tumour evolution during targeted therapy. Targeted therapy induces adaptations favoring APOBEC mutagenesis by inducing APOBEC3B in NF-κB dependent manner and UNG2 depletion due to c-Jun downregulation (genetic adaptation arising under therapy indicated by colored tumor cells in the resistant tumor).

We also identified a deleterious APOBEC signature mutation in *RMB10* (Q595*), which has been shown to diminish response to EGFR inhibitor treatment by providing protection from apoptosis, in a tumour intrinsically resistant to EGFR-targeted therapy^45^. The collective clinical case data suggest that the *APOBEC* upregulation (with *UNG* downregulation) that is induced by targeted therapy promotes the *de novo* genetic evolution of targeted therapy resistance via the acquisition of diverse ABOBEC-associated gene mutations.

## Discussion

Our study reveals that targeted therapy induces adaptations favoring APOBEC3B-mediated tumour genetic evolution (Fig. 4c). Our findings establish a solution for how tumour cells may acquire new mutations that are not present before treatment, and in particular in response to treatment, that contribute to tumour cell survival and thus the development of drug resistance. While APOBEC has been implicated in drug resistance previously^46,47^, our study reveals a distinct principle by which the cancer therapy actively drives upregulation of APOBEC to fuel the evolution of drug resistance: thus, the targeted cancer therapy drives tumour genetic evolution via APOBEC DNA deaminase engagement.

We demonstrated that expression of *APOBEC3B* drives resistance in pre-clinical cellular and mouse models of lung adenocarcinoma. Although we focused on oncogenic *EGFR*-driven lung adenocarcinomas, we also show that our findings extend to other molecular subsets as well such as *EML4-ALK*-driven lung cancer which suggests this may be a more general principle of therapy-induced genetic evolution. Our analysis of patient data also suggests an increase in *APOBEC3B* expression and APOBEC mutational signature in the post-TKI setting. We also identified specific APOBEC signature mutations that can drive resistance to targeted therapy through various mechanisms. Our data suggest that inhibiting APOBEC3B-driven tumour evolution could suppress the emergence of one key class of resistance mutations and thereby improve response to targeted therapy. Our work is consistent with and expands upon prior studies suggesting a potential association between APOBEC-mediated mutational signature and acquisition of putative resistance mutations in the APOBEC-preferred context during treatment of EGFR- and ALK-driven lung cancers that acquired resistance to targeted therapy^48,49^.

Our mouse models suggest that APOBEC3B-driven genetic adaptation is specific to a TKI therapeutic context, as APOBEC3B expression is detrimental at tumour initiation. In the TRACERx 100 dataset we observed an enrichment of the APOBEC mutation signature subclonally (Extended Data Fig 3)^10^. In addition, in a recent study of East Asian lung adenocarcinoma patients with a higher number of patients with EGFR driver mutations, the proportion of APOBEC-associated mutations was also elevated among late mutations^9^. Our data and these other patient-centered studies reinforce the idea that subclonal APOBEC genome mutagenesis may be selected for later during tumour evolution and in response to targeted therapy.

A previous study suggested a role for another APOBEC family member, AID, in the generation of *EGFR* T790M mutation during erlotinib treatment^50^. However, AID transcripts were undetectable in PC9 cells in our RNA Seq data. Furthermore, it is possible that other APOBEC3 family members beyond APOBEC3B might contribute to genetic evolution of resistance as well, an area for future investigation. Additionally, recent work showed that decreased mismatch repair capacity and increased expression of error prone DNA polymerases may contribute to increased adaptability of colorectal cancer to EGFR inhibition therapy^51^. While we observed downregulation of *UNG*, we did not observe upregulation of error prone polymerases in our RNA Seq data from EGFR TKI-treated *EGFR*-mutant PC9 NSCLC cells. Thus, our findings centering on APOBEC engagement describe a distinct mode and mechanism of therapy-induced genetic adaptability in cancer. Our evidence here and these emerging collective findings suggest that treatment-induced genetic mutability may be a general principle by which tumours evolve in response to treatment to become drug resistant, perhaps through context-specific or tumour-specific cellular mechanisms. Co-targeting APOBEC3 deaminases or other therapy-induced factors that promote this genetic evolution along with a primary driver oncoprotein such as oncogenic EGFR may be a promising strategy to suppress the emergence of drug resistance in cancer.

## Author contributions

**Conception and design of the study**: M.K.M., T.G.B, D.R.C., C.S., E.K.L., R.S.H., J.D.; **Data acquisition (cell line and animal studies)**: M.K.M, D.R.C., F.H., B.G., S.N., C.G., P.G., E.G., W.H., A.R., B.A., R.I.V., M.M., N.J.T., N.K., S.V., S.H.; **Data acquisition for clinical studies**: C.E.M., C.M.B., D.L.K., J.K.R., E.A.Y., L.T.; **Mutational Signature Analysis and/or other computational analysis**: N.I.V., N.A.T., W.W., L.C., E.M.V.A., J.Y.; **Analysis and interpretation of data**: M.K.M, D.R.C, N.I.V., T.G.B., C.S., E.K.L, R.S.H., W.L.B, L.K.L., C.D., P.P.A., J.P., M.A.B., A.N., M.D., C.M.R., S.K.C., N.M., C.M., B.R., B.B., W.W.; **Manuscript writing and revision**: M.K.M, D.R.C, T.G.B, C.S, E.M.V.A, R.S.H.

## Acknowledgements

This project is supported by the NIH/NCI U54CA224081, R01CA169338, R01CA211052, R01CA204302, U01CA217882 (to T.G.B). Pfizer, as well as the University of California Cancer League (United States) (to C.E.M), AstraZeneca (United Kingdom), The Damon Runyon Cancer Research Foundation P0528804 (United States), Doris Duke Charitable Foundation P2018110 (United States), V Foundation P0530519 (United States), and NIH/NCI R01CA227807 (to C.M.B.), F.H. was supported by the Mildred Scheel postdoctoral fellowship from the German Cancer Aid. E.A.Y is supported by T32 HL007185 from the NHLBI. Cancer studies in the Harris lab are supported in part by the National Cancer Institute P01-CA234228. R.S.H. is the Margaret Harvey Schering Land Grant Chair for Cancer Research, a Distinguished University McKnight Professor, and an Investigator of the Howard Hughes Medical Institute.

D.R.C was supported by the Francis Crick Institute which receives its core funding form Cancer Research UK (FC001169), the UK Medical Research Council (FC002269), and the Wellcome Trust (FC001169), as well as an NC3Rs training fellowship (NC/S001832/1). C.S. acknowledges grant support from Pfizer, AstraZeneca, Bristol Myers Squibb, Roche-Ventana, Boehringer-Ingelheim, Archer Dx Inc (collaboration in minimal residual disease sequencing technologies) and Ono Pharmaceutical, is an AstraZeneca Advisory Board member and Chief Investigator for the MeRmaiD1 clinical trial, has consulted for Pfizer, Novartis, GlaxoSmithKline, MSD, Bristol Myers Squibb, Celgene, AstraZeneca, Illumina, Genentech, Roche-Ventana, GRAIL, Medicxi, Bicycle Therapeutics, and the Sarah Cannon Research Institute, has stock options in Apogen Biotechnologies, Epic Bioscience, GRAIL, and has stock options and is co-founder of Achilles Therapeutics. R.S.H is a co-founder, shareholder, and consultant of ApoGen Biotechnologies Inc. The other University of Minnesota authors have no competing interests to declare. T.G.B. is an advisor to Novartis, Astrazeneca, Revolution Medicines, Array/Pfizer, Springworks, Strategia, Relay, Jazz, Rain, EcoR1 and receives research funding from Novartis and Revolution Medicines and Strategia. N.I.V. served on an Advisory Board for Sanofi Genzyme. E.M.V.A. is a consultant for Tango Therapeutics, Genome Medical, Invitae, Enara Bio, Janssen, Manifold Bio, Monte Rosa; receives research funding from Novartis, BMS; has equity in Tango Therapeutics, Genome Medical, Syapse, Enara Bio, Manifold Bio, Microsoft, Monte Rosa; has received travel reimbursement from Roche/Genentech and own institutional patents filed on chromatin mutations and immunotherapy response, and methods for clinical interpretation. C.E.M. is on advisory board of Genentech; receives honoraria from Novartis, Guardant, Research and receives funding from Novartis, Revolution Medicines. C.M.B. is a consultant for Amgen, Foundation Medicine, Blueprint Medicines, Revolution Medicines; receives research funding from Novartis, AstraZeneca, Takeda; and institutional research funding from Mirati, Spectrum, MedImmune and Roche.

Journal Editorial Board membership: C.S. is editorial board member for *Cell* and *PLOS medicine*, associate editor and editorial board member for *Annals of Oncology*, scientific editor for *Cancer Discovery* and advisory board member for *Nature Reviews Clinical Oncology* and *Cancer Cell*. T.G.B is editor-in-chief of *npj Precision Oncology* and an editorial board member of *Molecular and Cellular Biology*.

Patents: C.S. holds European patents relating to assay technology to detect tumour recurrence (PCT/GB2017/053289); to targeting neoantigens (PCT/EP2016/059401), identifying patent response to immune checkpoint blockade (PCT/EP2016/071471), determining HLA LOH (PCT/GB2018/052004), predicting survival rates of patients with cancer (PCT/GB2020/050221), identifying patients who respond to cancer treatment (PCT/GB2018/051912), a US patent relating to detecting tumour mutations (PCT/US2017/28013) and both a European and US patent related to identifying insertion/deletion mutation targets (PCT/GB2018/051892).

C.S. is Royal Society Napier Research Professor (RP150154). His work is supported by the Francis Crick Institute, which receives its core funding from Cancer Research UK (FC001169), the UK Medical Research Council (FC001169), and the Wellcome Trust (FC001169). C.S. is funded by Cancer Research UK (TRACERx, PEACE and CRUK Cancer Immunotherapy Catalyst Network), Cancer Research UK Lung Cancer Centre of Excellence, the Rosetrees Trust, Butterfield and Stoneygate Trusts, NovoNordisk Foundation (ID16584), Royal Society Research Professorship Enhancement Award (RP/EA/180007), the NIHR BRC at University College London Hospitals, the CRUK-UCL Centre, Experimental Cancer Medicine Centre and the Breast Cancer Research Foundation, USA (BCRF). His research is supported by a Stand Up To Cancer-LUNGevity-American Lung Association Lung Cancer Interception Dream Team Translational Research Grant (SU2C-AACR-DT23-17). Stand Up To Cancer is a program of the Entertainment Industry Foundation. Research grants are administered by the American Association for Cancer Research, the Scientific Partner of SU2C. C.S. also receives funding from the European Research Council (ERC) under the European Union’s Seventh Framework Programme (FP7/2007-2013) Consolidator Grant (FP7-THESEUS-617844), and ERC Advanced Grant (PROTEUS) from the European Research Council under the European Union’s Horizon 2020 research and innovation program (835297) and Chromavision from the European Union’s Horizon 2020 research and innovation programme (665233).

Special thanks to the Biological Research Facility at the Francis Crick Institute, specifically to Ade Adekoya, James Cormack, Antony Horwood, and Scott Lighterness for their hard work and support. Special thanks also to the Experimental Histopathology Laboratory at the Francis Crick Institute, specifically to Emma Nye, Bruna Almeida, Mary Green, and Richard Stone for their help and support. Special thanks to all the members of the Bivona laboratory (former and current), Dmitry Gordenin, Alejandro Sweet-Cordero, Sourav Bandyopadhyay, Marcus Breese, Swati Kaushik, Brandon Leonard and Sharath Raju for their insights and support and Sarah Elmes, Ashley Maynard, David V. Allegakoen and Abhik Tambe for their technical support.

## Methods

### Cell line and growth assays

All cell lines were grown in RPMI-1640 medium supplemented with 1% penicillin-streptomycin solution (10,000 Units/ml) and 10% FBS in a humidified incubator with 5% CO2 maintained at 37°C and fresh media was added to cells every 3-4 days. All drugs used for treatment except PBS-1086^17^ were purchased from Selleck Chemicals. For growth assays, cells were exposed to the indicated drugs for few days-2.5 months (long-term growth assays) or DMSO for 3-4 days (controls used for long-term growth assays) in 6 well plates or 96 well plates (details in the figure legends) and assayed using crystal violet staining or Celltiter-Glo luminescent viability assay (Promega) according to manufacturer’s instructions.

### Deriving clonal populations and generating APOBEC3B knockout cells

Clonal cells were derived by sorting single cells into 96 well plates and expanding them over a few weeks. We then derived pools of one of the clones expressing either a non-targeting guide or APOBEC3B targeting guide along and puromycin marker and CRISPR/Cas9 by lentiviral transduction as done in a previously published study^52^. A couple of gRNA target sequences, which were designed by the Zhang lab to specifically target APOBEC3B^53^ were first subcloned into the all-in-one lentiCRISPR v2 plasmid (Addgene plasmid # 52961- a gift from Feng Zhang)^53^. They were then lentivirally transduced into PC9 clonal cells, tested and the one that showed better APOBEC3B-depletion in western blot analysis was selected for further analysis. Additionally, we also generated APOBEC3B knockout clones from clonal derivatives of PC9 cells (C2 and E4). For this we used the Dox-inducible version of LentiCRISPR v2 (Addgene plasmid # 87360 - a gift from Adam Karpf) and subcloned the APOBEC3B-targeting sgRNA that worked well, into it^54^. The plasmid was then lentivirally transduced into PC9 clonal cells, subjected to puromycin selection and then half the cells were treated with 1 ug/ml doxycycline (dox) to induce the expression of Cas9 and thereby generate APOBEC3B knockouts. Induction of APOBEC3B deletion was confirmed by western blot analysis. Clonal cells were then derived from these cells using single cell sorting as indicated earlier and the clonal populations that deleted APOBEC3B were identified by western blot analysis and tested further. The cells not treated with dox served as APOBEC3B-proficient control for the APOBEC3B-knockout clones.

### Transductions and transfections

Hek293T cells were co-transfected with lentiviral packaging plasmids pCMVdr8 and pMD2.G plasmid, along with the plasmid of interest using Fugene 6 transfection reagent (Promega). All shRNA plasmids used were purchased from Sigma. Cells were transduced by incubation with 1:1 diluted lentivirus for 1-2 days and then selected with the antibiotic marker (puromycin or hyrgomycin) until untransduced control plate was clear. Negative control siRNA and other siRNA cocktails used were purchased from GE Dharmacon. siRNAs were transfected using Lipofectamine RNAi Max according to manufacturer’s protocol and the cells were harvested 48 hr of transfection for subsequent assays. RT-qPCR or western blot analysis were used to confirm the shRNA, siRNA of CRISPR-mediated depletions.

### RT-Qpcr

Total RNA was extracted using GeneJet RNA purification kit (Thermo Fisher) or RNeasy Mini kit (Qiagen) and cDNA was synthesized from it using sensiFast cDNA synthesis kit in accordance with the manufacturer’s instructions. qPCR reactions were performed using PowerUP SYBR Green Master Mix (Applied Biosystems) and previously validated primers^55^ (PrimerBank)} on a QuantStudio. GAPDH, 18S RNA or β2Microglobulin were used as reference genes. Data was analyzed using QuantStudio 12K Flex Software V1.3 and GraphPad Prism 7.

### Western Blot Assay

Cells were treated with the indicated drugs the day after plating on a 6 well plate or 10 cm plates for the indicated duration. Whole cell extracts were made with the former were harvested by first washing with ice cold PBS and then lysing using ice cold RIPA buffer containing protease and phosphatase inhibitors followed by sonication and centrifugation for clarification of extracts. Cells grown on 10 cm plates were used for making nuclear-cytoplasmic extracts as described previously using ice-cold 0.1% NP-40 in PBS^56^. Equal amounts of extracts (quantified using Lowry assay) were subjected to SDS-PAGE analysis on 4-15% Criterion TX gels (Bio-Rad), transferred to nitrocellulose membranes using semi-dry transfer apparatus and Tans-Blot Turbo RTA Midi Nitrocellulose Transfer kit (Bio-Rad). The membranes were then blocked with 3% milk in TBST, probed with primary antibody overnight at 4°C and then with the corresponding secondary antibody-either HRP-conjugated or fluorescently labelled for 1-2 hr at room temperature. They were then imaged on a Li-Cor imager or ImageQuant LAS 4000 (GE Healthcare). anti-APOBEC3B (5210-87-13)^57^ and anti-UNG^15^ antibodies were kindly provided by Reuben Harris and anti-GAPDH (sc-59540) was purchased from Santa Cruz Biotechnology. All other primary antibodies including anti-EGFR (#4267), anti-phospho-EGFR (Y1068, #3777 or #2236), anti-STAT3 (#9139), anti-phospho-STAT3 (Y705, #9145), anti-AKT (#2920), anti-phospho-AKT (S473, #4060), anti-phospho-ERK (T202, Y204; #4370 or #9106), anti-ERK (#9102), anti-RELA (#8242), anti-RELB (#4922), anti-Hsp90 (#4874), anti-TUBB (#2146) and anti-histone H3 (#9715) were purchased from Cell Signaling Technology.

### Enzymatic assays

APOBEC assays or Uracil Excision assays were performed by incubating nuclear extracts made using REAP method ^59^ with either of the following DNA oligo substrates (IDT): 5’ – ATTATTATTATTCAAATGGATTTATTTATTTATTTATTTATTT-FAM-3’ or 5’-AGCAGTATTUGTTGTCACGA-FAM-3’, respectively using established protocols^14,15^. Upon completion of the reactions, they were heated at 95 degrees for 5 minutes after addition of TBE Urea buffer (Novex) and immediately run on a 15% TBE-Urea gel (Bio-Rad). The gels were then imaged using Cy2 filter on ImageQuant LAS 4000 (GE Healthcare) and quantified using ImageJ and Microsoft Excel or GraphPad Prism 7.

### Subcutaneous Tumour Xenografts and PDX Studies

All animal experiments were conducted under UCSF IACUC-approved animal protocols. PC-9 tumour xenografts were generated by injection of one million cells in a 50/50 mixture for matrigel and PBS into 6- to 8-wk-old female NOD/SCID mice. Once the tumours grew to ∼200 mm^3^, the mice were then treated with vehicle or 5 mg/kg osimertinib once daily for 4 days and the tumours were harvested on day 4 and subsequently analyzed by western blot analysis. For H1975 xenografts, 1 million cells were injected into SCID mice and the indicated treatment were initiated (once daily, oral gavage) when the tumours were about 100 mm^3^. The tumour tissues were harvested on day 4 of treatment. PDX was generated as indicated in a previous study^17^. Tumours were passaged in SCID mice and treatment was initiated after the tumours were about 400 mm^3^. Tumours were harvested after 2 days of treatment with 25 mg/kg erlotinib, administered once daily by oral gavage.

### Mouse strains and tumour induction

The Cre-inducible *Rosa26::LSL-APOBEC3Bi* mice are described in the companion study (de Carné Trécesson et al 2020, manuscript under consideration). The *TetO-EGFR*^*L858R*^; *Rosa26*^*LNL-tTA*^ *(E)* and *CCSP-rtTA;TetO-EGFR*^*L858R*^*;Rosa26*^*CreER(T2)*^ mice have been described^19,20,58,59^. Tumours were initiated in *E* and *EA3B* mice by intratracheal infection of mice with adenoviral vectors expressing Cre recombinase as described^60^. Adenoviral-Cre (Ad-Cre-GFP) was from the University of Iowa Gene Transfer Core. Tumours were initiated in *EA3Bi* mice using chow containing doxycycline (625 ppm) obtained from Harlan-Tekland. All animal regulated procedures were approved by the Francis Crick Institute BRF Strategic Oversight Committee that incorporates the Animal Welfare and Ethical Review Body and conformed with the UK Home Office guidelines and regulations under the Animals (Scientific Procedures) Act 1986 including Amendment Regulations 2012.

### *In vivo* treatment with Erlotinib

Erlotinib was purchased from Selleckchem (Erlotinib Osi-744), dissolved in 0.3% methylcellulose and administered intraperitoneally at 25 mg/kg, 5 days a week.

### *In vivo* treatment with tamoxifen

Tamoxifen was administered by oral gavage 3 times in one week with at a 2-4 day intervals (3 injections total). Mice received tamoxifen at 150 mg/kg dissolved in sunflower oil.

### MicroCT Imaging

Mice were anaesthetized with isoflurane/oxygen for no more than an hour each. They were minimally restrained whilst anaesthetized during the actual imaging (usually 8-10 minutes). After scanning mice were observed and if necessary, placed in cages in a recovery chamber/rack until they regained consciousness and start to eat/drink. Tumour burden in each animal was quantified by calculating the tumour volume of visible tumours using AnalyzeDirect.

### Histological preparation and immunohistochemical staining

Tissues were fixed in 10% formalin overnight and transferred to 70% ethanol until paraffin embedding. IHC was performed using the following primary antibodies: EGFR^L858R^ mutant specific (Cell Signaling: 3197, 43B2), APOBEC3B (5210-87-13)^57^, Ki67 (Abcam: Ab15580), Caspase 3 (R&D (Bio-Techne): AF835), p-Histone H2AX (Sigma-Aldrich, Ref. no. 05-636) and UNG (NB600-1031, Novus Biologicals). Sections were developed with DAB and counterstained with hematoxylin. The number of EGFR^L858R^, APOBEC3B, Ki67, Caspase 3, and gH2AX positive cells were quantified using QuPath.

### Tumour processing

Tumour tissue was collected form -80° freezer and placed on dry ice. Tissue was placed on a petri dish and cut into pieces. RLT Buffer with β-Mercaptoethanol was dispensed into 15 ml reaction vessels. A small part of the tumour was added to the lysis buffer, and remaining parts were refrozen. TissueRuptor was used for disruption and homogenization of tissue. The tip of the disposable probe was plced into the vessel and the TissueRuptor was turned to speed 1 until the lysate was homogeneous (usually about 10 seconds). Lysate was added to a previously prepared QIAshredder tube and centrifuged at full speed for 1 minute. The homogenised solution was then added to AllPrep DNA spin columns. The Qiagen AllPrep DNA/RNA Mini Kit (80204).

### Cell line Whole Genome Mutational Signature Analysis

Sequences were aligned to the human genome (hg38) using the Burrows-Wheeler Aligner (version 0.7.17). PCR duplicates were removed using Picard (version 2.18.16). Reads were locally realigned around InDels using GATK3 (version 3.6.0) tools RealignerTargetCreator to create intervals, followed by IndelRealigner on the aligned bam files. MuTect2 from GATK3 (version 3.6.0) was used in tumour/normal mode to call mutations in test vs control cell lines. Single nucleotide variants (SNVs) that passed the internal GATK3 filter with read depths over 30 reads at called positions, at least 4 reads in the alternate mutation call and an allele frequency greater than 0.1 were used for downstream analysis. Figures were plotted using the deconstructSigs R package^61^.

### DNA and RNA isolation from cell line models for sequencing

DNA or RNA were extracted using from frozen cell pellets using Qiagen’s DNeasy Blood and tissue kit or Qiagen’s RNeasy mini kit, respectively, as per the manufacturer’s instructions. The isolated DNA or RNA was quantified and qualitatively assessed using Qubit Fluorometer (Thermo Fisher) and a Bioanalyzer (Agilent) as per the manufacturer’s instructions. The DNA or RNA were then sent to BGI for whole genome sequencing (30X) or Novogene for mRNA or whole exome sequencing.

### Human Subjects

All patients gave informed consent for collection of clinical correlates, tissue collection, research testing under Institutional Review Board (IRB)-approved protocols (CC13-6512 and CC17-658, NCT03433469). Patient demographics are listed in Supplemental Tables. Patient studies were conducted according to the Declaration of Helsinki, the Belmont Report, and the U.S. Common Rule.”

### Studies with specimens from lung cancer patients

Frozen or formalin-fixed paraffin embedded (FFPE) tissues from lung cancer patients for DNA or RNA Sequencing (Bulk and Single Cell) studies were, processed and sequenced as described previously^18,42^. Some of these biopsies were subjected to whole exome sequencing at the QB3-Berkley Genomics for which library preparation was performed using IDT’s xGen exome panel. For additional specimens, tumour DNA from formalin-fixed paraffin embedded (FFPE) tissues and matched non-tumour from blood aliquots or stored buffy coats were collected as part of University of California, San Francisco’s (UCSF) biospecimen resource program (BIOS) in accordance with UCSF’s institutional review board approved protocol. DNA from blood aliquots were isolated at the BIOS. Other non-tumour samples and FFPE tumour tissues were sent for extraction, assessment of quality and quantity to Novogene and those meeting the required samples standards were subjected to whole exome sequencing at Novogene’s sequencing facility.

### Mutation analysis

Paired-end reads were aligned to the hg19 human genome using the Picard pipeline (https://gatk.broadinstitute.org/). A modified version of the Broad Institute Getz Lab CGA WES Characterization pipeline (https://docs-google-com.ezp-prod1.hul.harvard.edu/document/d/1VO2kX_fgfUd0=3mBS9NjLUWGZu794WbTepBel3cBg08) was used to call, filter and annotate somatic mutations. Specifically, single-nucleotide variants (SNVs) and other substitutions were called with MuTect (v1.1.6)^62^. Mutations were annotated using Oncotator^63^. MuTect mutation calls were filtered for 8-OxoG artifacts, and artifacts introduced through the formalin fixation process (FFPE) of tumour tissues ^67^. Indels were called with Strelka (v1.0.11). MuTect calls and Strelka calls were further filtered through a panel of normal samples (PoN) to remove artifacts generated by rare error modes and miscalled germline alterations^62^. To pass quality control, samples were required to have <5% cross-sample contamination as assessed with ContEst^62^; mean target coverage of at least 25x in the tumour sample and 20x in the corresponding normal as assessed using GATK3.7 DepthOfCoverage; and a percentage of tumour-in-normal of < 30% as determined by deTiN^64^. This pipeline was modified for analysis of cell lines rather than tumour-normal pairs as follows: indels were called through MuTect2 alone rather than Strelka; deTiN was not performed; and a common variant filter was applied to exclude variants present in The Exome Aggregation Consortium (ExAC) if at least 10 alleles containing the variant were present across any subpopulation, unless they appeared in a list of known somatic sites^65,66^.

### Mutational signature analysis

Active mutational processes^36^ were determined using the *deconstructSigs* R package^64^, with a signature contribution cutoff of 6%. This cutoff was chosen because it was the minimum contribution value required to obtain a false-positive rate of 0.1% and false-negative rate of 1.4% per the authors’ in-silico analysis, and is the recommended cutoff^61^. Samples with < 10 mutations were excluded from analysis due to poor signature discrimination with so few mutations and a sample with less than 15 days of exposure to TKI therapy was excluded because it is too short a time to accumulate detectable mutations due to therapy. For TRACERx data analysis data processing were performed in the R statistical environment version > = 3.3.1.

### RNA-Seq analyses

PDX-tissue RNA extractions were carried using RNeasy micro kit (Qiagen). RNA-Seq was performed using replicate samples on the Illumina HiSeq4000, paired-end 100-bp reads at the Center for Advanced Technology (UCSF). For the differential gene expression analysis, DESeq program was used to compare controls to erlotinib samples as previously described^67^.

RNAseq samples from patients and cell lines were sequenced by Novogene (https://en.novogene.com/) with paired-end sequencing (150bp in length). There were ∼20 million reads for each sample. The processed fastq files were mapped to hg19 reference genome using STAR (version 2.4) algorithm and transcript expressions were quantified using RSEM (version 1.2.29) algorithm. The default parameters in the algorithms were used. The normalized transcript reads (TPM) were used for downstream analysis.

For single cell RNA Seq analyses the data from a previously published study (all cancer cells from advanced lung cancer patients) was used and analyzed in a similar manner^18^. All cells used are identified as malignant by marker expression and CNV inference and originated in from various biopsy sites (adrenal, liver, lymph node, lung, pleura/pleural fluid). Nonparametric, pairwise comparisons (Wilcoxon Rank Sum Test) was used to determine the statistical significance of the pairwise comparisons of different timepoints for their average scaled expression.

### Statistical Analysis

One-way or two-way ANOVA test with Holm’s Sidak correction for multiple comparisons (>2 groups) or two-tailed t-test with Welch’s correction (2 groups) were used to determine the statistical significance of the differences between groups for RT-qPCR, growth and enzymatic assays and Bulk RNA Seq analysis. Normality of immunohistochemical and microCT data was determined using multiple testing methods (Anderson-Darling test, D’Agostino & Pearson test, Shapiro-Wilk test, and Kolmogorov-Smirnov test). A two-sided t-test or two-sided Mann-Whitney test was used for immunohistochemical and microCT data depending on the normality tests to determine the statistical significance of the differences between groups. Analysis for these assays was done using Graphpad Prism 7.

## Data availability

Data used in Ext. Fig. 5a were downloaded from Sequence Read Archive with accession number SRP068321^34^. Data used in Ext. Fig. 5b were downloaded from Sequence Read Archive with accession number SRP022943^33^. For single cell RNA Seq analyses shown in Ext. Fig. 4g-i the data from a previously published study (all advanced lung cancer cell data) were used and analyzed in a similar manner^18^. This data is available as an NCBI Bioproject # PRJNA591860. The RNA Seq data for Ext. Fig.4a was from a previously published study^17^. These data are available at NCBI GEO under accession number GSE65420.

---

**Extended Fig. 1.**
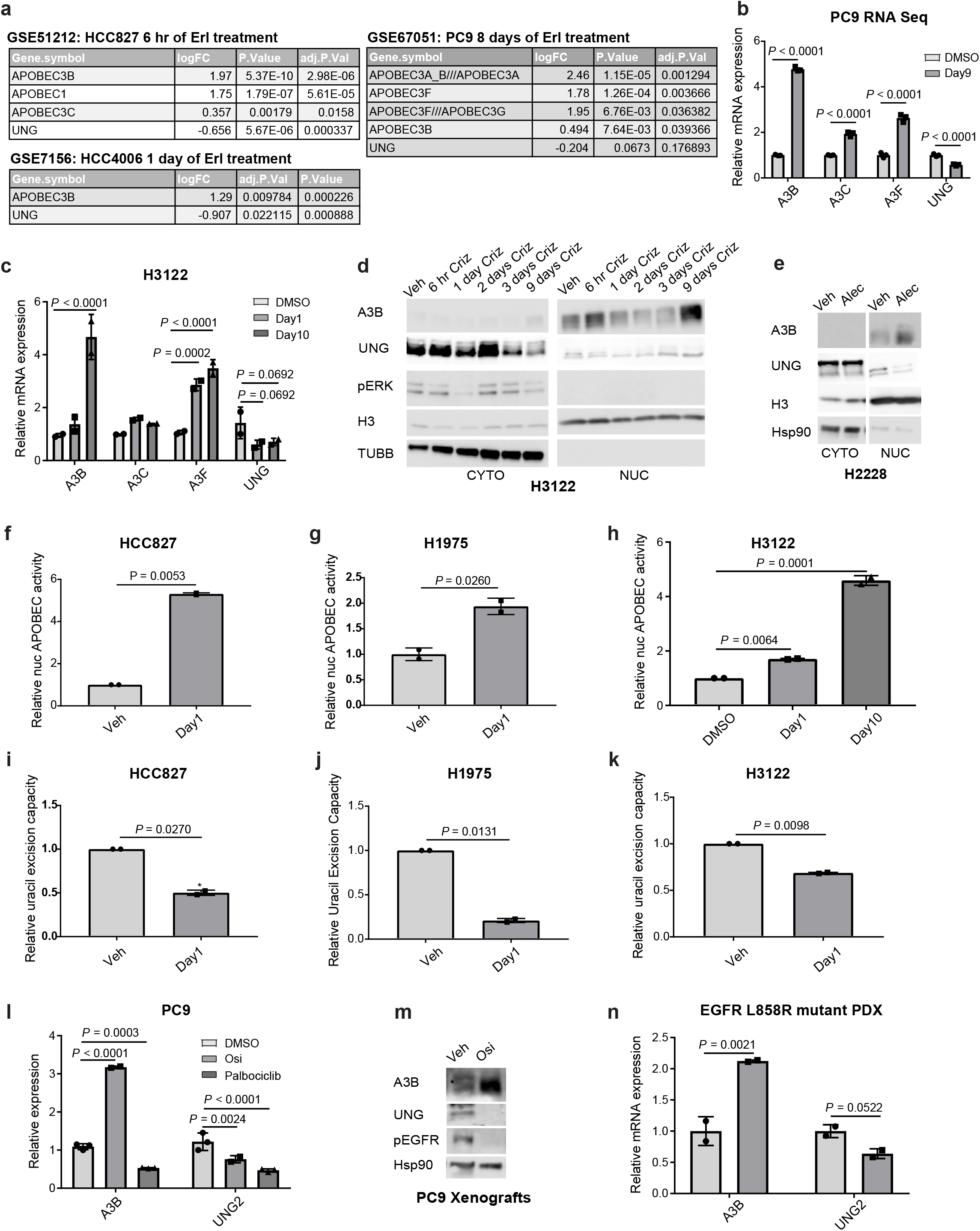
Treatment with tryosine kinase inhibitors (TKIs) induces adaptations conducive to APOBEC-mediated mutagenesis in multiple pre-clinical models of lung adenocarcinoma. **a**, GEO2R analysis of the indicated GSEA datasets of *EGFR*-driven cellular models of human lung adenocarcinoma treated with erlotinib (erl). **b**, Gene expression changes in PC9 cells under treatment (2 μM osimertinib for 9 days) relative to DMSO treated cells identified using RNA Seq analysis (n = 3 biological replicates, mean ± SEM, ANOVA test). **c**, RT-qPCR analysis of H3122 cells after 18 hours of treatment with DMSO or 0.5 μM crizotinib or 10 days of treatment with 0.5 μM crizotinib (mean ± SD, n = 2 biological replicates, ANOVA test). **d**, Western blot analysis of extracts from EML4-ALK positive H3122 human NSCLC cells treated with DMSO or 1 μM crizotinib (criz) for the indicated durations. **e**, Western blot analysis of EML4-ALK positive H2228 human NSCLC cells treated with DMSO or 0.5 μM alectinib (alec) for 18 hours (CYTO: cytoplasmic extracts; NUC: nuclear extracts). **f-k**, APOBEC activity assay (**f-h**) and uracil excision capacity assay (**i-k**) performed using nuclear extracts of *EGFR*-mutant HCC827 or H1975 NSCLC cells treated with DMSO versus 0.4 μM osimertinib or DMSO versus 1 μM osimertinib, respectively for 18 hours or H3122 cells treated with DMSO versus 0.5 μM crizotinib for 18 hours and/or 10 days (n = 2 biological replicates, mean ± SD, ANOVA or t-test). **l**, RT-qPCR analysis of PC9 cells treated with DMSO, 2 μM osimertinib (osi) of 1 μM of Palbociclib for 18 hours (n = 2 or 3 biological replicates, mean ± SD, ANOVA test). **m**, Western blot analysis of extracts of PC9 tumour xenografts treated with vehicle or 5 mg/kg osimertinib. **n**, Gene expression analysis using RNA seq analysis upon treatment of a PDX model of human *EGFR*-driven lung adenocarcinoma with vehicle or erlotinib (two biological replicates, mean ± SD, ANOVA test). (H3: Histone H3).

**Extended Fig. 2.**
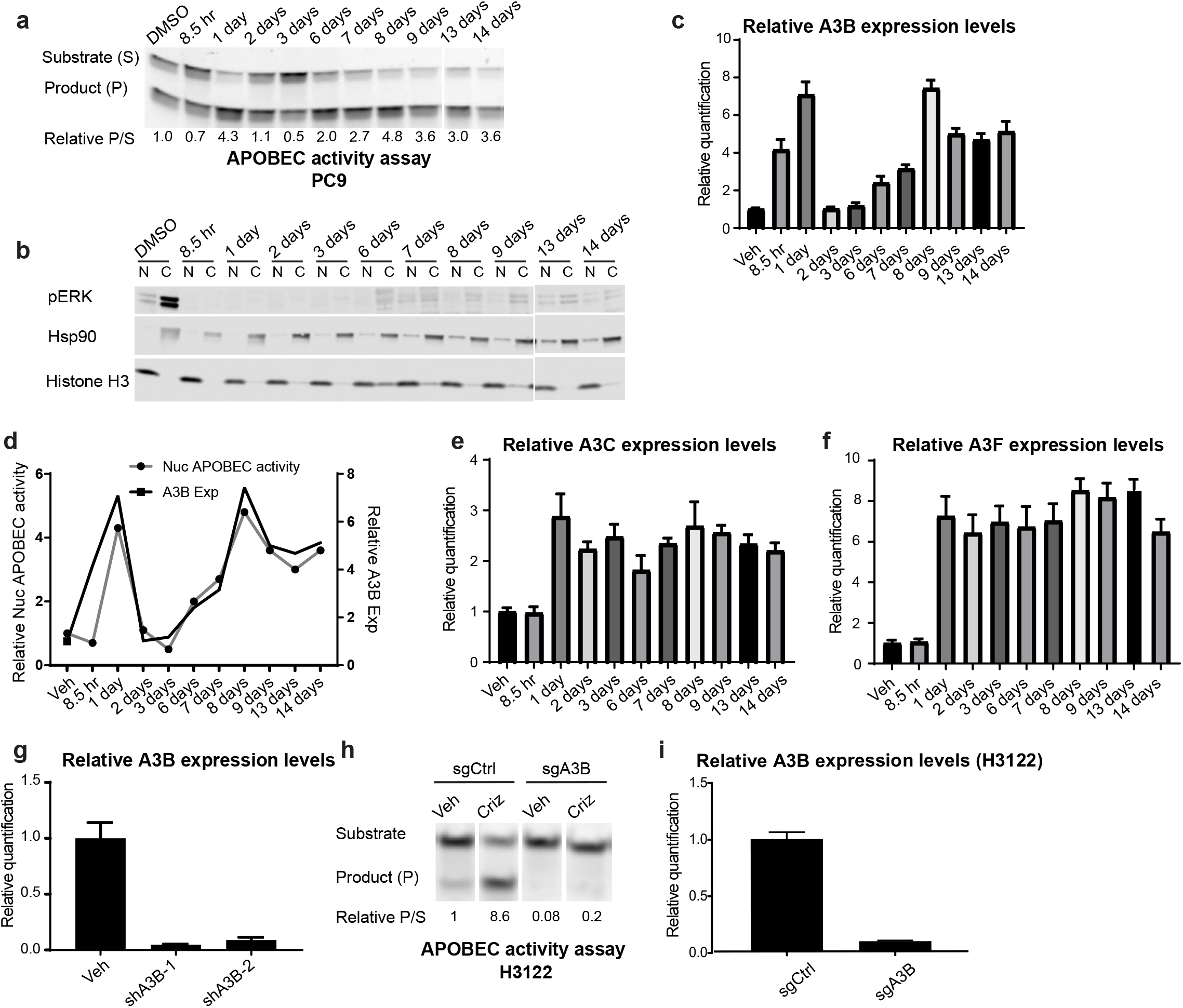
APOBEC3B drives a TKI-induced increase in nuclear APOBEC activity. **a, b**, APOBEC activity assay performed using nuclear extracts of PC9 cells treated with increasing doses of osimertinib (200 nM-up to 3 days, 3 to 6 days-1 μM osimertinib, 6 to 14 days 2 μM osimertinib) and western blot analysis of these extracts (C: Cytoplasmic extract; N: Nuclear extract). **c, e-f**, RT-qPCR-based examination of the expression of the indicated APOBEC3 family members in PC9 cells treated similar to that as Ext. Fig. **2a** (assayed in triplicate, mean ± SD). **d**, Comparison of the pattern of changes in nuclear APOBEC activity and APOBEC3B (A3B) expression after osimertinib treatment shown in Ext. Fig. **2a** and **2b. g, i**, RT-qPCR-based validation of APOBEC3B knockdown in PC9 cells (assayed in triplicate, mean ± SD). **h**, APOBEC activity assay using nuclear extracts of H3122 cells transduced with derivatives of lentiCRISPR v2 encoding non-targeting (sgCtrl) or APOBEC3B targeting (sgA3B) sgRNA. **i**, RT-qPCR analysis of H3122 cells (assayed in triplicate, mean ± SD) grown together with cells used for nuclear APOBEC activity assay in **h**.

**Extended Fig. 3.**
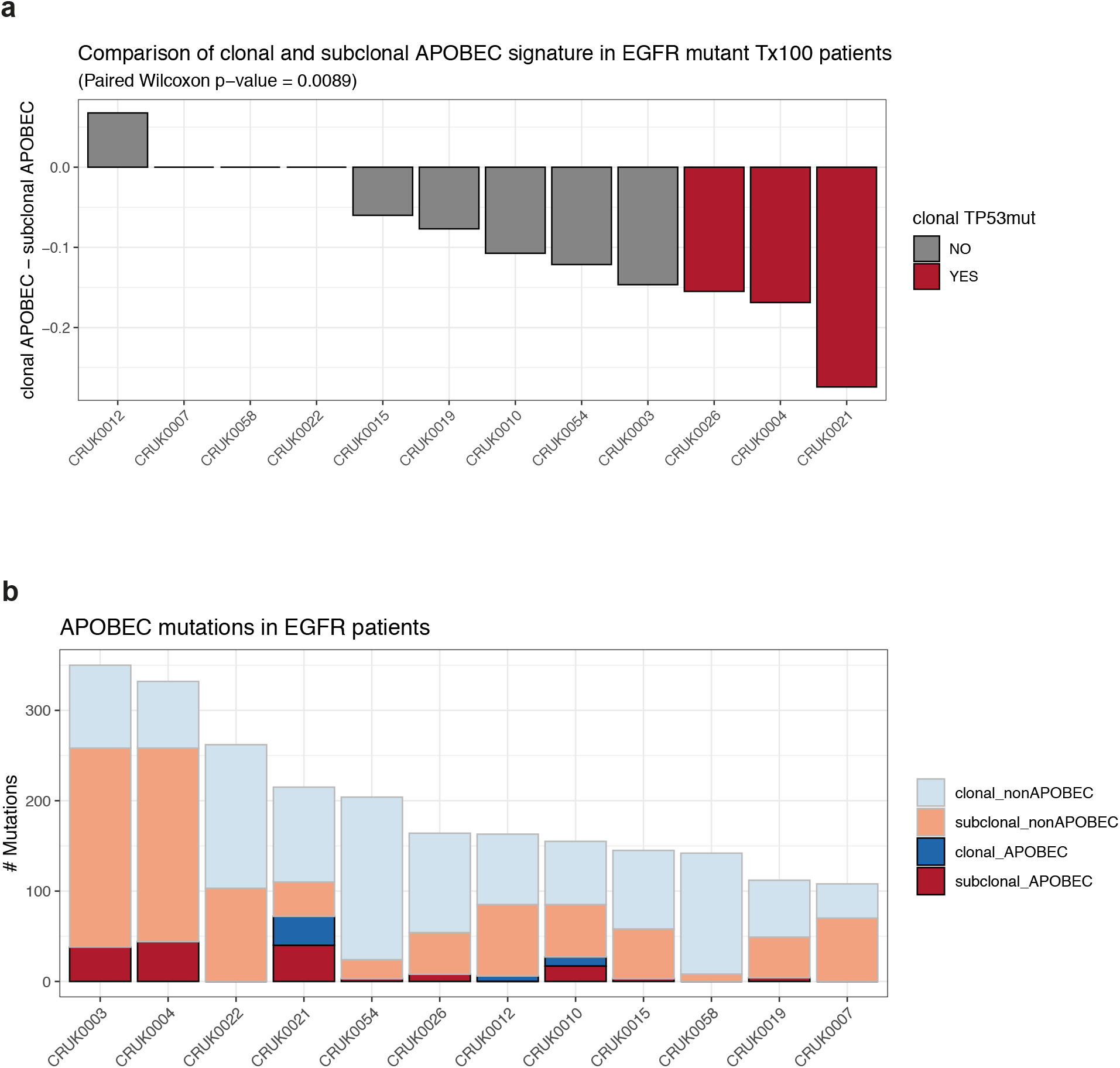
Subclonal APOBEC mutagenesis in EGFR mutant patients from TRACERx 100 dataset. **a**, Comparison of clonal and subclonal APOBEC mutation signature (clonal APOBEC - subclonal APOBEC) in patients with EGFR driver mutations (1, 1a, exon 19 deletion). Grey bars indicate the patient is TP53 wildtype or has a subclonal TP53 mutation. Red bars indicate that the patient has a clonal TP53 mutation. P< 0.01. **b**, Number of APOBEC mutations in patients with EGFR driver mutations (1, 1a, exon 19 deletion). Colors indicate clonal or subclonal APOBEC or non APOBEC mutations.

**Extended Fig. 4.**
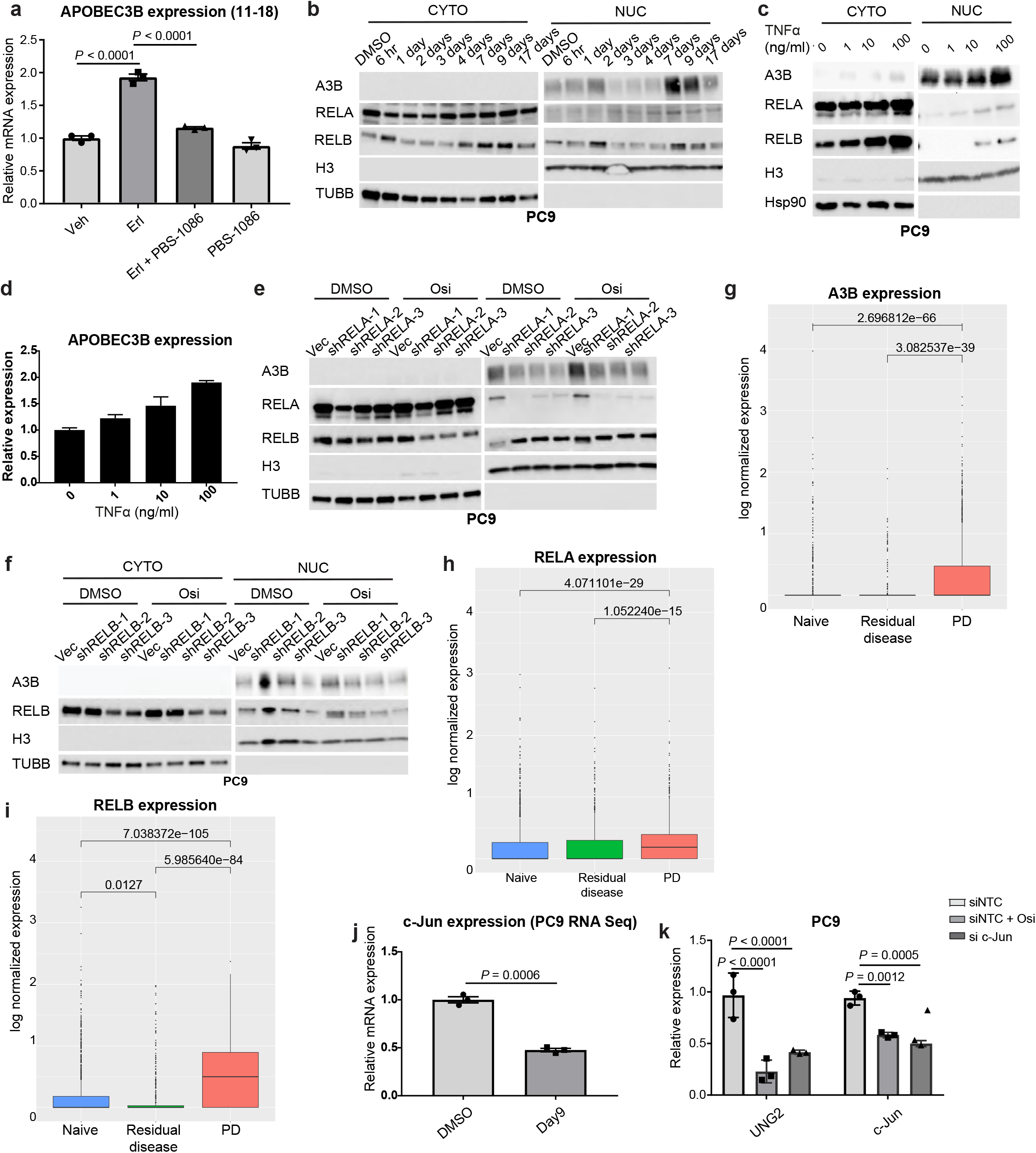
Mechanisms inducing adaptations favoring APOBEC-mutagenesis after TKI treatment. **a**, RNA Seq analysis *EGFR*-mutant 11-18 NSCLC cells treated with DMSO, 100 erlotinib or 5 μM PBS-1086 (NF-κB inhibitor) individually or in combination (n = 3 biological replicates, mean ± SEM, ANOVA test). **b, c, e, f** Western blot analysis of extracts from PC9 cells treated with DMSO or 4 μM osimertinib (osi) for the indicated durations (**b**), or treated with TNFα for 8.5 hours (hrs) (**c**), or transduced with vector or plasmids encoding shRNA targeting RELA (**e**) or RELB (**f**) and treated with DMSO or 0.2 μM osimertinib for 18 hours (CYTO: cytoplasmic extracts; NUC: nuclear extracts). **d**, RT-qPCR analysis of TNFα-treated PC9 cells assayed in triplicate. **g-i** Box plots showing single cell RNA Seq analysis of lung cancer cells from patient tumors across different treatment time points (Wilcoxon Rank Sum test). **j**, Gene expression analysis from RNA Seq experiment with osimertinib-treated NSCLC cells (PC9) shown in Fig. **1b. k**, RT-qPCR analysis of PC9 cells transfected with non-targeting siRNA (siNTC) or c-Jun targeting siRNA and treated with DMSO or 2 μM osimertinib for 18 hours (n = 3 biological replicates; mean ± SD, ANOVA test). (H3: Histone H3, TUBB: Tubulin Beta Class I).

**Extended Fig. 5.**
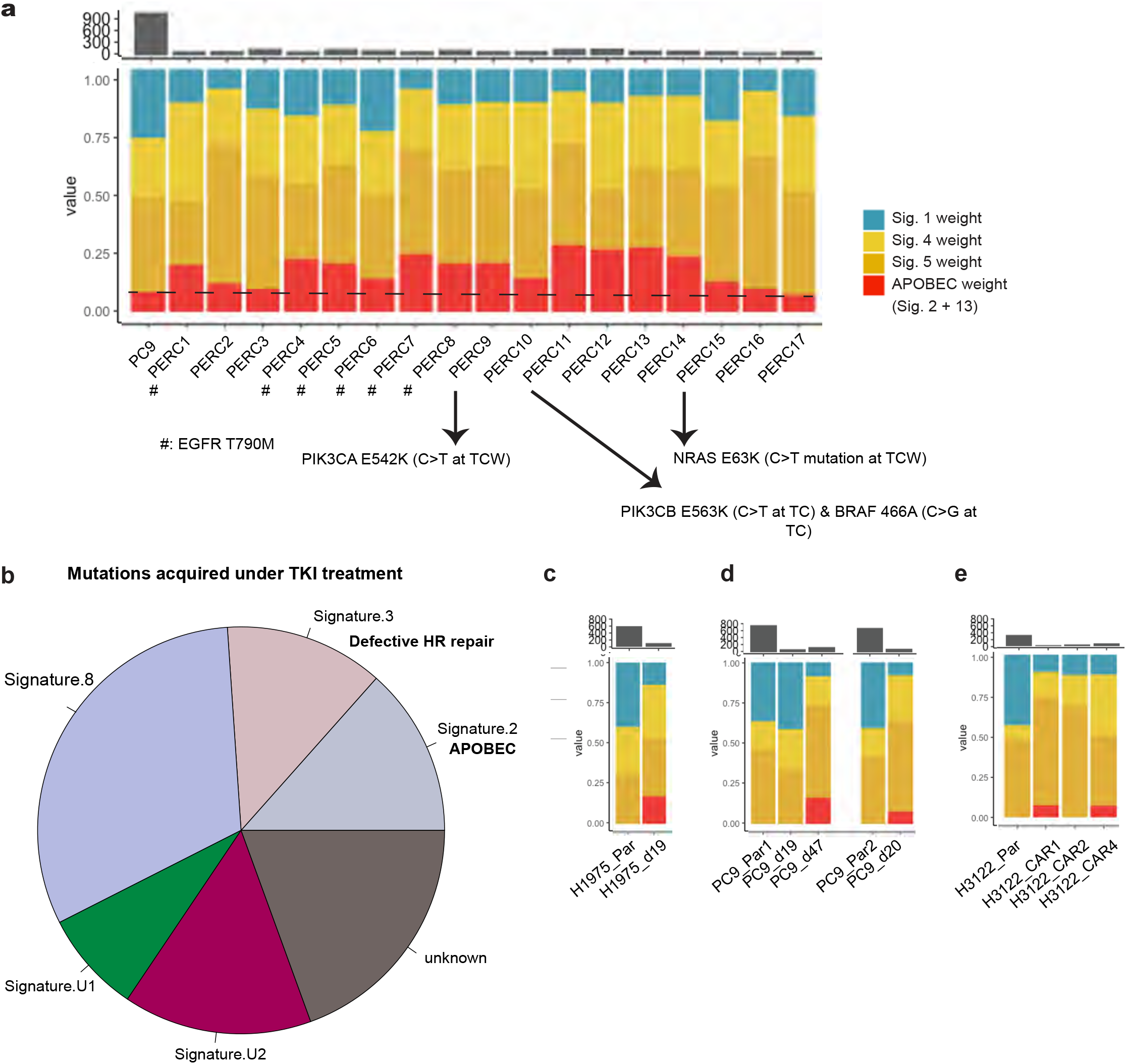
Evidence for acquisition of APOBEC-favored mutagenesis and mechanisms of resistance in pre-clinical models of lung adenocarcinoma. **a**, Mutational signature and additional mutational analysis of mutations present in PC9 parental cells and mutations acquired by PC9 clonal resistant cells during long term treatment with erlotinib (PERC: Persister derived erlotinib resistant clones) ^35^. **b**, Mutational signature analysis of mutations acquired during erlotinib treatment versus those present in the vehicle exposed *EGFR*-mutant NSCLC cells, derived by comparing whole genome sequencing data from parental and resistant cell line derivatives which harbor the *EGFR* T790M mutation ^36^. The proposed aetiology (if known) is indicated below the signature in bold (HR: homologous recombination) ^38^. **c, d**, Mutational signature analysis of mutations in the indicated vehicle treated parental cells and mutations detected after indicated duration with osimertinib treatment (Par: parental cells). **e**, Mutational signature analysis of mutations in parental cells or acquired during increasing doses of crizotinib treatment used for deriving the resistant cells {(CAR: crizotinib acquired resistance) ^45^ (Par: parental cells)}. (The axis label ‘value’ represents the relative proportions of the indicated mutational mutational signatures).

**Extended Fig. 6.**
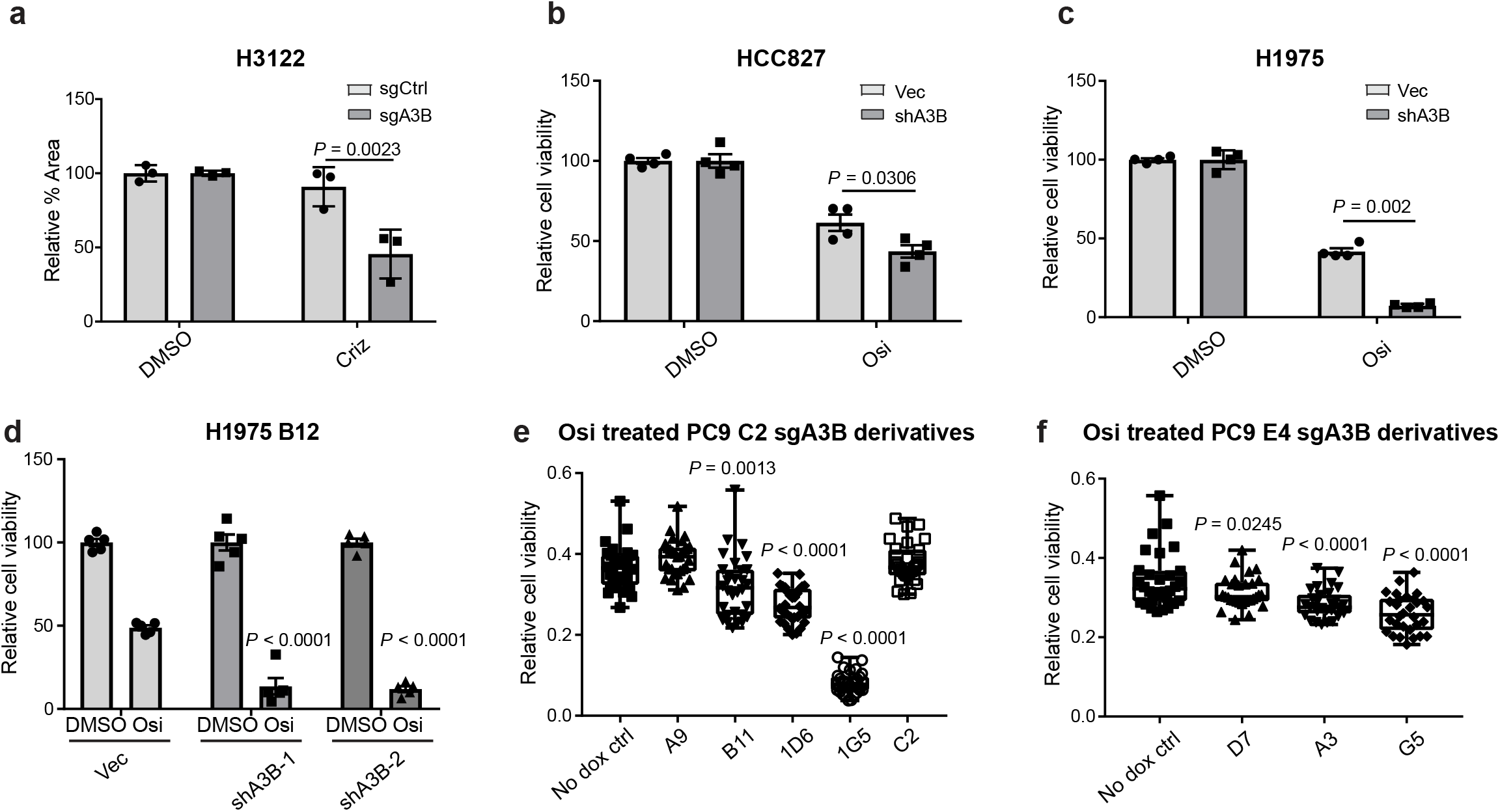
Depletion of APOBEC3B impairs survival of lung adenocarcinoma cells subjected to TKI treatment. **a**, Crystal violet growth assays (area of crystal violet staining in the indicated wells relative to their DMSO controls) comparing long-term growth of APOBEC3B-proficient and APOBEC3B-deficient cells in the presence of crizotinib (criz) (n = 3 biological replicates, mean ± SD, t-test, 100 nM - 8 days, 50 nM - 3 days, 100 nM - 4 days, 1-2 uM Criz - 17 days). **b-d**, Cell-titre glo viability assays performed on the indicated APOBEC3B-deficient or APOBEC3B-proficient cells subjected to high dose treatment initially followed by low dose treatment over a long-term with the osimertinib (osi) (n=4 or 5 biological replicates, mean ± SD, ANOVA/t-test). ({HCC827: osi treatment:: 2 uM-first 5 days, 100 nM - 22 days from day6 onwards}, {H1975: osi treatment-4 μ M Osi - 8 days}, {H1975 B12: osi treatment-2 μM - 7 days, low dose i.e. 12.5-50 nM - over the next 15 days). **e, f**, Cell-titre go viability assays performed on derivatives of PC9 C2 and E4 clonal cells transduced with pTLCV2 plasmid derivative expressing sgRNA targeting APOBEC3B without or with three-day Cas9 induction with 1 μg/ml doxycycline to induce APOBEC3B knockout. Signal in long-term osimertinib treated wells normalized to the signal from DMSO treated cells. (n = 30, whiskers: min to max, ANOVA test), comparing each of the validated clonal APOBEC3B(A3B)-knockout derivatives (A9, B11, 1D6, 1G5, C2, D7, A3, G5) with the no doxycycline control (osi treatment duration: 2.5 months total; dose: 100 nM - first 4 days, high dose (500 nM-2 μM) - 42 days from day5 onwards, 100 nM - remaining duration).

**Extended Fig. 7.**
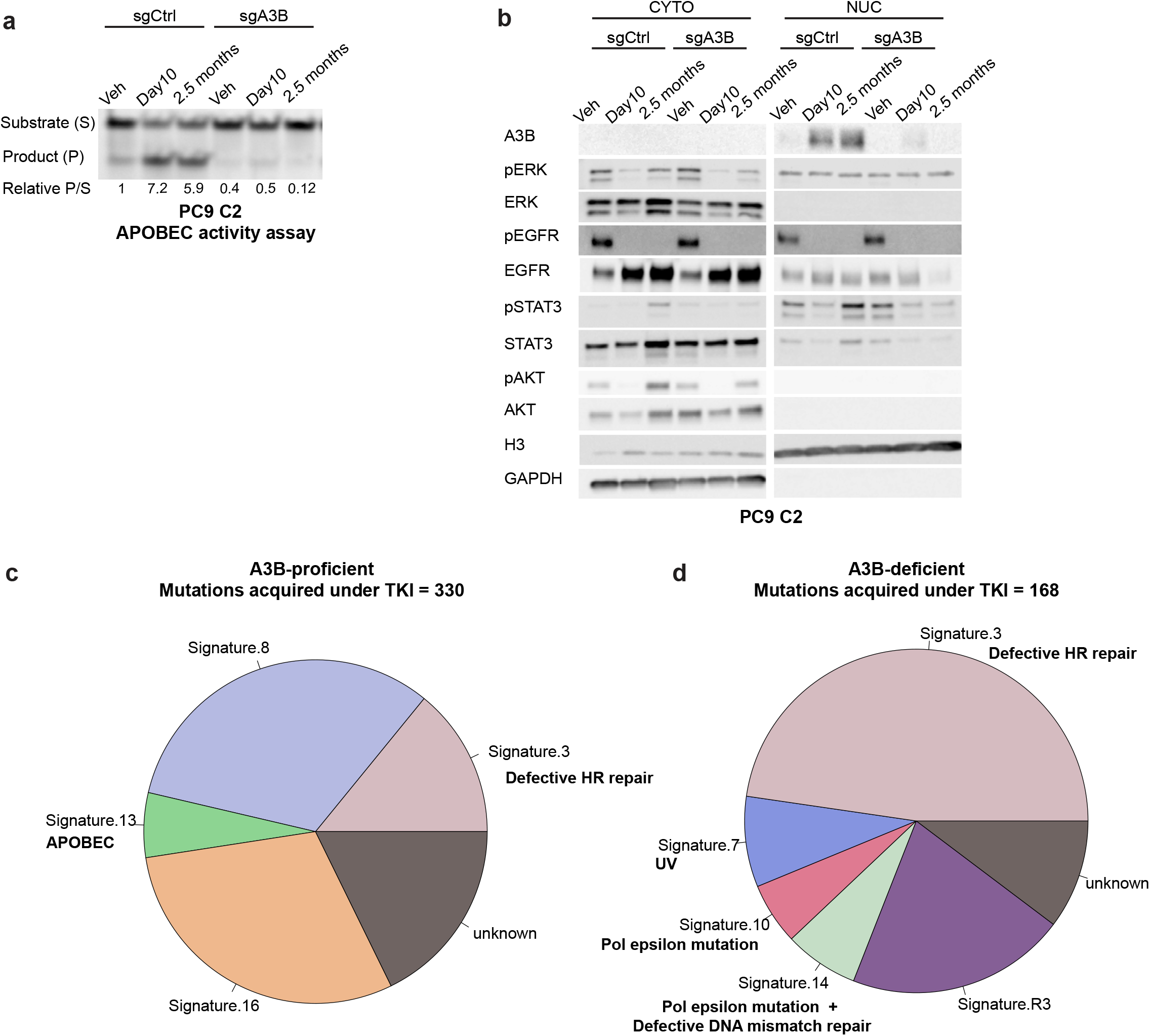
APOBEC3B-dependent evolution during TKI treatment. **a, b**, Nuclear APOBEC activity assay and western blot analysis of clonal PC9 cells (C2) transduced with lentiCRISPRV2 plasmid derivatives encoding non-targeting (sgCtrl) or APOBEC3B-targeting (sgA3B) sgRNA and treated with DMSO (Veh-Vehicle) or osimertinib for the indicated duration (CYTO - cytoplasmic extracts, NUC - nuclear extracts). **c, d**, Mutational signature analysis of mutations acquired during indicated durations of osimertinib treatment performed using whole genome sequencing data derived from DNA extracted from DMSO treated and cells treated with osimertinib shown in **a** (osi treatment duration: 2.5 months; dosing: 100 nM-first 4 days, high dose (1 μM-2 μM) - 52 days from day5 onwards, subsequently 100 nM for 18 days and 25 nM for 1 day) The proposed aetiology (if known) is indicated below each signature in bold (HR: homologous recombination; Pol: DNA polymerase) ^38^.

**Extended Fig. 8.**
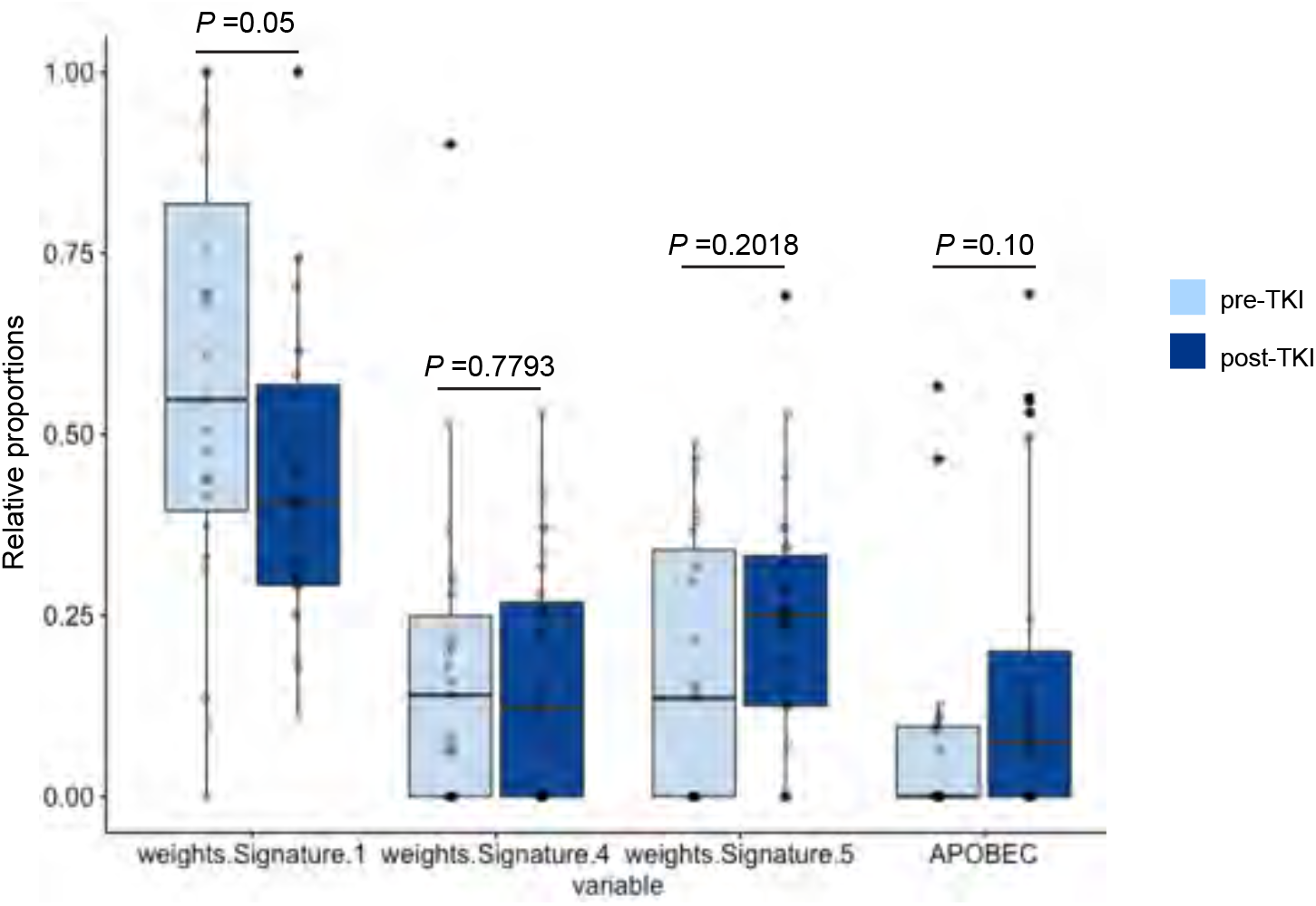
Comparion of proportions of mutational signatures in human lung cancers pre- and post-TKI therapy. Analyses comparing the proportion of the most commonly observed mutational signatures (Signature 1,4, 5 and APOBEC) in patients prior to and after TKI therapy (Wilcoxon test).

