## Supplementary Tables 1-3 for "Targeted cancer therapy induces APOBEC fuelling the evolution of drug resistance"

**Supplementary Table 1:** Demographic information of the patients and their tumour specimens biopsied for bulk RNA Seq analysis.

| patient_id | biopsy_site | driver_mutation | gender | race | smoking hx | Classifier | treatment_history |
|---|---|---|---|---|---|---|---|
| TH217 | LN | EGFR del19 | M | Asian | Never | Treatment naive | tx naive |
| TH218 | Adrenal | EGFR L858R | F | Asian | Never | Treatment naive | tx naive |
| TH223 | LN | EGFR del19 | F | White or Caucasian | Never | Treatment naive | tx naive |
| TH225 | LN | KRAS G12C | F | White or Caucasian | Former | Treatment naive | tx naive |
| TH227 | LN | ALK intron 19 rearrangement | M | White or Caucasian | Never | Treatment naive | tx naive |
| TH226 | Lung | EGFR del19 | M | White or Caucasian | Former | Treatment naive | tx naive |
| TH231 | Lung | ALK fusion | M | White or Caucasian | Never | Treatment naive | tx naive |
| TH_179_E2_T1a | lung | BRAF V600E | M | white | Never | Treatment naive | tx naive |
| TH_187_E3_T1 | lung | MET exon 14 splice | F | asian | Never | Treatment naive | tx naive |
| TH_169_E2_T1 | lung | EGFR Exon 19 del | F | asian | Never | Treatment naive | tx naive |
| TH_208_E1_T1 | LN | EGFR L861Q | F | white | Never | Treatment naive | tx naive |
| TH_210_E1_T1 | abd mass | ALK | M | black | Never | Treatment naive | tx naive |
| TH_218_E4_T1 | adrenal | EGFR L858R | F | Asian | Never | Treatment naive | tx naive |
| TH_226_E2_T1 | lung | EGFR Exon 19 del | M | white | Former | Treatment naive | tx naive |
| TH_231_E1_T1 | LN | ALK-EML4 | M | white | Never | Treatment naive | tx naive |
| TH_231_E2_T1A | Lung | ALK-EML4 | M | white | Never | Treatment naive | tx naive |
| TH_238_E1_T1A | lung | BRAF V600E | F | white | Former | Treatment naive | tx naive |
| TH_205_E2_T1 | pleura | EGFR L858R | M | asian | Never | Treatment naive | tx naive |
| TH_067_E4_T1a | liver | EGFR Exon 19 del | F | white | Never | Treatment naive | tx naive |
| AZ05SCRN | lung | EGFR L858R | F | asian | Never | Treatment naive | tx naive |
| AZ003SCRN | lung | EGFR L858R | F | white | Former | Treatment naive | tx naive |
| AZ009SCRN | lung | EGFR Exon 19 del | F | asian | Never | Treatment naive | tx naive |
| AZ010SCRN | lung | EGFR Exon 19 del | M | white | Former | Treatment naive | tx naive |
| TH205 | Lung | EGFR L858R | M | Asian | Never | Residual disease | on firsline TKI |
| TH218 | Adrenal | EGFR L858R | F | Asian | Never | Residual disease | on firsline TKI |
| TH226 | Lung | EGFR del19 | M | White or Caucasian | Former | Residual disease | on treatment |
| TH_067_E7_Liver Met k | liver (autopsy) | EGFR Exon 19 del | F | white | Never | Residual disease | chemo/ TKI/IO |
| TH_107_E4_T1 | lung | EGFR Exon 19 del | M | black | Never | Residual disease | TKI |
| TH_158_E7_T1 | lung | EGFR Exon 19 del | F | Pacific Islander | Never | Residual disease | TKI |
| TH_169_E4_T1a | lung | EGFR Exon 19 del | F | asian | Never | Residual disease | TKI |
| TH_171_E2_T1 | LN | EML4-ALK fusion Variant 3a/b | F | asian | Never | Residual disease | chemo/ tki/io |
| TH_178_E3_T1 | LN | CD74-ROS1 | F | white | Never | Residual disease | chemo/ tki/io |
| TH_205_E3_T1a | lung | EGFR L858R | M | asian | Never | Residual disease | none |
| TH_208_E4_pl cells 1 | pleural fluid | EGFR L861Q | F | white | Never | Residual disease | none |
| TH_218_E6_T1a | adrenal | EGFR L858R | F | Asian | Never | Residual disease | none |
| TH_226_E3_T1a | lung | EGFR Exon 19 del | M | white | Former | Residual disease | none |
| TH_187_E5_T1a | lung | MET exon 14 splice | F | asian | Never | Residual disease | none |
| TH_067_E5_T1 | liver | EGFR Exon 19 del | F | white | Never | Residual disease | none |
| TH_067_E6_T1a | liver | EGFR Exon 19 del | F | white | Never | Residual disease | chemo/ tki/io |
| TH_187_E6_T1A | lung | MET exon 14 splice | F | asian | Never | Residual disease | none |
| AZ003SURG | lung | EGFR L858R | F | white | Former | Residual disease | none |
| AZ005SURG | lung | EGFR L858R | F | asian | Never | Residual disease | none |
| AZ008SURG | lung | EGFR Exon 19 del | F | white | Former | Residual disease | none |
| AZ010SURG | lung | EGFR Exon 19 del | M | white | Former | Residual disease | none |
| TH220 | Liver | ALK fusion | F | White or Caucasian | Never | Progressive disease | Prior TKI/ immunotherapy |
| TH155 | Brain | EGFR del19 | F | Native Hawaiian or Other Pacific Island | Never | Progressive disease | on firstline TKI |
| TH067 | Liver | EGFR del19 | F | White or Caucasian | Never | Progressive disease | Prior chemo/ TKI/IO |
| TH222 | LN | ROS1 fusion | F | Asian | Never | Progressive disease | on treatment |
| TH_116_E2_T1a | liver | EGFR L858R | F | asian | Never | Progressive disease | TKI |
| TH_208_E3_asp_ppt 1 | lung/ pleural fluid | EGFR L861Q | F | white | Never | Progressive disease | none |
| TH_041_E2_T1 | ovary | EML4-ALK V3a/b | F | white | Former | Progressive disease | chemo/TKI |
| TH_107_E5_T1a | lung | EGFR Exon 19 del | M | black | Never | Progressive disease | TKI |
| TH_155_E5_T1 | brain | EGFR Exon 19 del | F | Pacific Islander | Never | Progressive disease | TKI |
| TH_169_E6_T1a | lung | EGFR Exon 19 del | F | asian | Never | Progressive disease | TKI |
| TH_171_E3_M1a(cv) | LN | EML4-ALK fusion Variant 3a/b | F | asian | Never | Progressive disease | chemo/ tki/io |

**Supplementary Table 2:** Demographic information of the patients and their tumour specimens biopsied for whole exome sequencing and subsequent mutational signature analysis and inclusion/exclusion criteria.

| participant | pair_id | case_sample | control_sample | ContEst Fraction | TiN | tumor_mean_ depth_of_cvg | normal_mean_ depth_of_cvg | Excluded | Exclusion_reason | Treatment_status_therapy | reported_ driver | detected_driver | driver_overall | TP53_ nonsyn | all_muts | all_SNVs |
|---|---|---|---|---|---|---|---|---|---|---|---|---|---|---|---|---|
| TH11 | TH11_pre_1_norm | TH11_pre_1 | TH11_Norm | 0.648 | NA | 1.53 | 80.12 | 1 | Multiple | NA | NA | NA | NA | NA | NA | NA |
| TH97 | TH97_a_norm | TH97_a | TH97_Norm | 0.728 | NA | 2.4 | 195.79 | 1 | Multiple | NA | NA | NA | NA | NA | NA | NA |
| TH84 | TH84_a1_norm | TH84_a1 | TH84_Norm | 0.322 | 0 | 2.44 | 95.12 | 1 | Low coverage_tumor | TKI | EGFR | none | EGFR | 0 | 74 | 42 |
| TH91 | TH91_c_norm | TH91_c | TH91_Norm | 0.513 | 0 | 2.95 | 162.61 | 1 | Multiple | TKI | EGFR | none | EGFR | 0 | 118 | 18 |
| TH11 | TH11_pre_2_norm | TH11_pre_2 | TH11_Norm | 0.135 | 0 | 3.84 | 80.12 | 1 | Multiple | Pre_treatment_2 | EGFR | none | EGFR | 0 | 292 | 173 |
| TH208 | TH208E3_WB | TH208E3 | TH208WB | NA | NA | 3.89 | 58.8 | 1 | Low coverage_tumor | NA | NA | NA | NA | NA | NA | NA |
| TH294 | TH294E1_WB | TH294E1 | TH294WB | 0.122 | 0 | 5.24 | 35.06 | 1 | Low coverage_tumor | Chemo | EGFR | none | EGFR | 0 | 105 | 63 |
| TH11 | TH11_post_1_norm | TH11_post_1 | TH11_Norm | 0.036 | 0 | 7.8 | 80.12 | 1 | Low coverage_tumor | TKI | EGFR | EGFRp.T790M | EGFR | 0 | 230 | 143 |
| TH80 | TH80_A1_norm | TH80_A1 | TH80_Norm | 0.341 | 0 | 8.44 | 146.77 | 1 | Multiple | Pre_treatment | EGFR | none | EGFR | 0 | 521 | 221 |
| TH80 | TH80_B2_norm | TH80_B2 | TH80_Norm | 0.147 | NA | 9.06 | 146.77 | 1 | Multiple | NA | NA | NA | NA | NA | NA | NA |
| TH84 | TH84_c2_norm | TH84_c2 | TH84_Norm | 0.05 | 0 | 9.3 | 95.12 | 1 | Low coverage_tumor | Pre_treatment | EGFR | EGFRp.L858R | EGFR | 0 | 167 | 75 |
| TH273 | | TH273E2 | | NA | NA | 14.87 | NA | 1 | Low coverage_tumor | NA | NA | NA | NA | NA | NA | NA |
| TH56 | TH56_2_norm | TH56_2 | TH56_Norm | 0.053 | 0 | 17.08 | 44 | 1 | Multiple | TKI | EGFR | none | EGFR | 0 | 16 | 15 |
| TH74 | TH74_a_norm | TH74_a | TH74_Norm | 0.043 | 0 | 22.86 | 120.8 | 1 | Low coverage_tumor | Pre_treatment | EGFR | EGFRp.L747_P753delinsS | EGFR | 1 | 820 | 201 |
| TH239 | TH239E4_WB | TH239E4 | TH239WB | 0.011 | 0 | 25.68 | 73.07 | NA | | TKI | EGFR | EGFRp.E709A, p.G719S | EGFR | 0 | 244 | 122 |
| TH29 | TH29_Post_norm | TH29_Post | TH29_Norm | 0.04 | 0 | 28.7 | 72.78 | NA | | TKI | EGFR | none | EGFR | 0 | 14 | 12 |
| TH239 | TH239E5_WB | TH239E5 | TH239WB | 0.02 | 0 | 28.98 | 73.07 | NA | | Pre_treatment | EGFR | none | EGFR | 0 | 70 | 15 |
| TH56 | TH56_c_norm | TH56_c | TH56_Norm | 0.04 | 0 | 31.87 | 44 | 1 | Mut_count_below10 | Pre_treatment | EGFR | none | EGFR | 0 | 10 | 9 |
| TH51 | TH51_A2_norm | TH51_A2 | TH51_Norm | 0.028 | 0 | 46.55 | 49.64 | NA | | TKI | EGFR | EGFRp.L858R | EGFR | 0 | 205 | 200 |
| TH17 | TH17_pre_norm | TH17_pre | TH17_Norm | 0.005 | 0 | 51.91 | 66.43 | NA | | Pre_treatment | EGFR | EGFRp.E746Ifs*16, p.R748Qfs*16 | EGFR | 0 | 43 | 40 |
| TH51 | TH51_B_norm | TH51_B | TH51_Norm | 0.08 | 0 | 52.35 | 49.64 | 1 | High_contest | Pre_treatment | EGFR | EGFRp.L858R | EGFR | 0 | 32 | 27 |
| CB2 | CB2_BX-2_norm | CB2_BX-2 | CB2_Norm | 0.02 | 0 | 58.38 | 96.41 | NA | | TKI | EGFR | EGFRp.L861Q, p.L62R | EGFR | 0 | 37 | 31 |
| TH79 | TH79_a_norm | TH79_a | TH79_Norm | 0.03 | 0 | 60.15 | 122.27 | NA | | Pre_treatment | EGFR | EGFRp.E746_A750del | EGFR | 1 | 271 | 53 |
| TH56 | TH56_B1_norm | TH56_B1 | TH56_Norm | 0.052 | 0 | 61.36 | 44 | 1 | High_contest | NA | NA | NA | NA | NA | NA | NA |
| TH56 | TH56_A1_norm | TH56_A1 | TH56_Norm | 0.024 | 0 | 62.37 | 44 | 1 | Mut_count_below10 | TKI | EGFR | none | EGFR | 0 | 6 | 5 |
| TH91 | TH91_a_norm | TH91_a | TH91_Norm | 0.047 | 0 | 71.18 | 162.61 | NA | | Pre_treatment | EGFR | EGFRp.L858R, p.R108G | EGFR | 0 | 68 | 61 |
| TH29 | TH29_Pre_norm | TH29_Pre | TH29_Norm | 0.008 | 0 | 71.26 | 72.78 | NA | | Pre_treatment | EGFR | EGFRp.L747_P753delinsS | EGFR | 0 | 61 | 58 |
| TH74 | TH74_2_norm | TH74_2 | TH74_Norm | 0.004 | 0 | 86.58 | 120.8 | NA | | TKI | EGFR | EGFRp.T790M, p.L747_P753delinsS | EGFR | 1 | 75 | 64 |
| TH194 | TH194E2_WB | TH194E2 | TH194WB | 0.031 | 0 | 93.76 | 63.5 | NA | | Chemo_TKI | EGFR | none | EGFR | 0 | 16 | 10 |
| TH11 | TH11_post_2_norm | TH11_post_2 | TH11_Norm | 0.008 | 0 | 98.41 | 80.12 | NA | | TKI | EGFR | EGFRp.E746_A750del, p.T790M | EGFR | 1 | 81 | 64 |
| TH087 | TH087E5_WB | TH087E5 | TH087WB | 0.017 | 0 | 98.44 | 53.88 | NA | | Chemo_TKI | EGFR | EGFRp.E746_A750del | EGFR | 1 | 574 | 559 |
| TH52 | TH52_A1_pre_norm | TH52_A1_pre | TH52_Norm | 0.002 | 0.05 | 100.5 | 145.49 | NA | | Pre_treatment | EGFR | EGFRp.L858R | EGFR | 0 | 72 | 61 |
| TH153 | TH153E3_WB | TH153E3 | TH153WB | 0.004 | 0 | 102.04 | 66.69 | NA | | TKI | EGFR | EGFRp.T790M | EGFR | 0 | 124 | 111 |
| TH087 | TH087E2_WB | TH087E2 | TH087WB | 0.019 | 0 | 103.67 | 53.88 | 1 | Mut_count_below10 | Chemo | EGFR | none | EGFR | 0 | 15 | 5 |
| CB2 | CB2_BX-1_norm | CB2_BX-1 | CB2_Norm | 0.007 | 0 | 112.43 | 96.41 | NA | | Pre_treatment | EGFR | EGFRp.L62R, p.L861Q | EGFR | 0 | 56 | 52 |
| TH97 | TH97_2_norm | TH97_2 | TH97_Norm | 0.006 | 0 | 116.73 | 195.79 | 1 | Mut_count_below10 | TKI | EGFR | none | EGFR | 0 | 4 | 4 |
| TH079 | TH079E2_WB | TH079E2 | TH079WB | 0.001 | 0 | 117.4 | 75.13 | NA | | TKI | EGFR | EGFRp.E746_A750del, p.T790M | EGFR | 1 | 320 | 300 |
| TH52 | TH52_C_post_norm | TH52_C_post | TH52_Norm | 0.005 | 0.02 | 128.96 | 145.49 | NA | | TKI | EGFR | EGFRp.L858R, p.T790M | EGFR | 0 | 124 | 113 |
| CB1 | CB1_BX-1_norm | CB1_BX-1 | CB1_Norm | 0.004 | 0.02 | 139.88 | 134.55 | NA | | Pre_treatment | EGFR | EGFRp.L858R, p.L62R | EGFR | 1 | 118 | 103 |
| Patient_13 | SGK_preRx_norm | SGK_preRx | SGK_Norm | 0.003 | 0 | 150.7 | 99.43 | NA | | Pre_treatment | EGFR | EGFRp.L858R | EGFR | 1 | 130 | 123 |
| AZ003 | AZ03SCRN_WB | AZ03SCRN | AZ03WB | 0.015 | 0 | 159.15 | 87.88 | NA | | Pre_treatment | none | EGFRp.L858R | EGFR | 1 | 139 | 135 |
| TH179 | TH179_E4_T1b_norm | TH179_E4_T1 | TH179_Norm | 0.001 | 0 | 172.42 | 96.68 | | | TKI | BRAF | none | BRAF | 0 | 14 | 14 |
| TH194 | TH194E1_WB | TH194E1 | TH194WB | 0.001 | 0 | 172.54 | 63.5 | NA | | Chemo | EGFR | EGFRp.T790M | EGFR | 0 | 303 | 276 |
| AZ009 | AZ09SURG_WB | AZ09SURG | AZ09WB | 0.001 | 0 | 172.65 | 72.37 | NA | | TKI | none | none | none | 1 | 97 | 93 |
| TH266 | TH-266-E2_norm | TH-266-E2 | TH266_Norm | 0.001 | 0.14 | 174.39 | 193.45 | | | Pre_treatment | none | none | none | 0 | 25 | 22 |
| TH155 | TH155_E5_norm | TH155_E5 | TH155_Norm | 0.001 | 0 | 174.94 | 73.24 | NA | | TKI | EGFR | EGFRp.V1109Ifs*6 | EGFR | 1 | 84 | 77 |
| TH187 | TH187_E3_norm | TH187_E3 | TH187_Norm | 0.001 | 0 | 178.44 | 102.48 | NA | | Pre_treatment | MET | METp.X1010_splice | MET | 1 | 172 | 164 |
| AZ003 | AZ03SURG_WB | AZ03SURG | AZ03WB | 0.001 | 0 | 181.27 | 87.88 | NA | | TKI | none | EGFRp.L858R | EGFR | 1 | 112 | 109 |
| TH148 | TH148E1_WB | TH148E1 | TH148WB | 0.001 | 0 | 181.82 | 23.21 | NA | | Pre_treatment | EGFR | none | EGFR | 1 | 114 | 109 |
| TH179 | TH179_E1_norm | TH179_E1 | TH179_Norm | 0.001 | 0 | 182.95 | 96.68 | NA | | TKI | BRAF | BRAFp.V640E | BRAF | 0 | 81 | 75 |
| TH256 | TH256E4_WB | TH256E4 | TH256WB | 0.001 | 0.07 | 188.35 | 61.41 | NA | | Pre_treatment | EGFR | EGFRp.L62R | EGFR | 0 | 45 | 45 |
| TH203 | TH203E4_WB | TH203E4 | TH203WB | 0.001 | 0 | 190.19 | 73.6 | NA | | Pre_treatment | EGFR | none | EGFR | 1 | 136 | 124 |
| TH169 | TH169_E2_norm | TH169_E2 | TH169_Norm | 0.001 | 0 | 194.17 | 93.14 | NA | | Pre_treatment | EGFR | none | EGFR | 0 | 93 | 84 |
| TH226 | TH226_E3_norm | TH226_E3 | TH226_Norm | 0.003 | 0 | 196.11 | 103.78 | NA | | TKI | EGFR | none | EGFR | 1 | 141 | 131 |
| TH067 | TH67_E4_norm1 | TH067_E4 | TH067_Norm_ | 0.001 | 0 | 202.43 | 276.6 | NA | | Pre_treatment | EGFR | none | EGFR | 0 | 197 | 183 |
| TH172 | TH172_E3_norm | TH172_E3 | TH172_Norm | 0.004 | 0 | 207.16 | 77.99 | | | TKI | BRAF | BRAFp.V640E | BRAF | 0 | 174 | 167 |
| TH231 | TH231_E1_norm | TH231_E1 | TH231_Norm | 0.001 | 0 | 210.94 | 100.85 | | | Pre_treatment | ALK | none | ALK | 0 | 47 | 46 |
| TH208 | TH208_E3_norm | TH208_E3 | TH208_Norm | 0.001 | 0 | 219.05 | 99.71 | NA | | TKI | EGFR | EGFRp.L861Q | EGFR | 1 | 130 | 120 |
| TH210 | TH210_E1_norm | TH210_E1 | TH210_Norm | 0.002 | 0 | 223.73 | 108.91 | NA | | Pre_treatment | ALK | none | ALK | 0 | 127 | 113 |
| TH205 | TH205_E2_norm | TH205_E2 | TH205_Norm | 0.001 | 0 | 226.04 | 121.48 | | | Pre_treatment | EGFR | none | EGFR | 1 | 69 | 66 |
| TH122 | TH122E3_WB | TH122E3 | TH122WB | 0.001 | 0 | 229.24 | 73.84 | NA | | TKI | EGFR | EGFRp.E746_A750del, p.T790M | EGFR | 0 | 145 | 130 |
| TH310 | TH310E3_WB | TH310E3 | TH310WB | 0.003 | 0 | 229.49 | 59.14 | NA | | Pre_treatment | EGFR | none | EGFR | 1 | 126 | 120 |
| TH116 | TH116_E2_norm | TH116_E2 | TH116_Norm | 0.001 | 0.01 | 229.81 | 99.12 | NA | | TKI | EGFR | EGFRp.L858R | EGFR | 1 | 125 | 122 |
| TH171 | TH171_E3_norm | TH171_E3 | TH171_Norm | 0.001 | 0 | 236 | 92.87 | NA | | Multiple_chemo_TKI | ALK | ALKp.I1322M, p.E1210K | ALK | 0 | 220 | 216 |
| TH169 | TH169_E4_norm | TH169_E4 | TH169_Norm | 0.001 | 0 | 239.73 | 93.14 | NA | | TKI | EGFR | none | EGFR | 0 | 116 | 106 |
| TH176 | TH176E2_WB | TH176E2 | TH176WB | 0.001 | 0 | 242.66 | 55.99 | NA | | Pre_treatment | EGFR | none | EGFR | 0 | 218 | 213 |
| AZ005 | AZ05SURG_WB | AZ05SURG | AZ05WB | 0.001 | 0 | 246.78 | 70.16 | | | TKI | none | EGFRp.L858R | EGFR | 0 | 97 | 97 |
| TH275 | TH275E1_WB | TH275E1 | TH275WB | 0.003 | 0 | 253.98 | 59.21 | NA | | Pre_treatment | EGFR | EGFRp.N158= | EGFR | 0 | 84 | 82 |
| TH330 | TH330E1_WB | TH330E1 | TH330WB | 0.001 | 0 | 256.02 | 75.11 | NA | | TKI | EGFR | none | EGFR | 1 | 148 | 141 |
| TH220 | TH220_E2_norm | TH220_E2 | TH220_Norm | 0.001 | 0 | 267.36 | 91.17 | NA | | Multiple_chemo_TKI | ALK | none | ALK | 0 | 142 | 133 |
| TH41 | TH41_E2_norm | TH041_E2 | TH041_Norm | 0.001 | 0.12 | 272.2 | 86.4 | | | Chemo/TKI | ALK | none | ALK | 0 | 109 | 107 |
| TH330 | TH330E2_WB | TH330E2 | TH330WB | 0.005 | 0 | 277.26 | 75.11 | NA | | Pre_treatment | EGFR | none | EGFR | 1 | 85 | 83 |
| AZ010 | AZ10SURG_WB | AZ10SURG | AZ10WB | 0.001 | 0 | 288.41 | 74.46 | NA | | TKI | none | none | none | 0 | 79 | 74 |
| TH208 | TH208_E4_norm | TH208_E4 | TH208_Norm | 0.001 | 0 | 338.23 | 99.71 | NA | | TKI | EGFR | EGFRp.L861Q | EGFR | 1 | 153 | 144 |
| TH067 | TH67_E7_norm1 | TH067_E7 | TH067_Norm_ | 0.001 | 0 | 338.9 | 276.6 | NA | | Multiple_chemo_TKI | EGFR | EGFRp.E746Vfs*16, p.L747Ffs*14 | EGFR | 0 | 430 | 412 |
| TH17 | TH17_post_norm | TH17_post | TH17_Norm | NA | NA | NA | 66.43 | 1 | Failed alignment | NA | NA | NA | NA | NA | NA | NA |
| Patient_13 | SGK_postRx_norm | SGK_postRx | SGK_Norm | NA | NA | NA | 99.43 | 1 | Failed alignment | NA | NA | NA | NA | NA | NA | NA |

**Supplementary Table 2:** Demographic information of the patients and their tumour specimens biopsied for whole exome sequencing and subsequent mutational signature analysis and inclusion/exclusion criteria.

| participant | Alias | pair_id | case_sample | control_sample | Clinical Oncogene | Treatment.status.at.biopsy | Treatment_status_condensed | Treatment_status_relative_to_prior_therapies | Sample Source | Treatment history | Smoking history | Reported_driver | detected_driver | driver_overall | TP53_nonsyn | all_muts | all_nonsyns | all_syns | all_SNVs | all_INS | all_DEL | all_DNP |
|---|---|---|---|---|---|---|---|---|---|---|---|---|---|---|---|---|---|---|---|---|---|---|
| AZ003 | Patient_1 | AZ03SCRN_WE | AZ03SCRN | AZ03WB | EGFR L858R | Pre Treatment | Pre_treatment | Pre_treatment | | N/A | | EGFR | EGFRp.L858R | EGFR | 1 | 139 | 84 | 55 | 135 | 0 | 4 | 0 |
| TH067 | Patient_2 | TH67_E4_norm1 | TH067_E4 | TH067_Norm_1 | EGFR Exon 19 del | treatment naive | Pre_treatment | Pre_treatment | liver | none | Never | EGFR | none | EGFR | 0 | 197 | 125 | 72 | 183 | 4 | 9 | 1 |
| TH79 | Patient_3 | TH79_a_norm | TH79_a | TH79_Norm | EGFR 746_750del | Pre Treatment | Pre_treatment | Pre_treatment | Primary tumor | N/A | | EGFR | EGFRp.E746_A750del | EGFR | 1 | 271 | 92 | 179 | 53 | 84 | 134 | 0 |
| TH330 | Patient_4 | TH330E2_WB | TH330E2 | TH330WB | EGFR L858R | Pre Treatment | Pre_treatment | Pre_treatment | No path | No Tx | Never | EGFR | none | EGFR | 1 | 85 | 54 | 31 | 83 | 0 | 1 | 1 |
| Patient_13 | Patient_5 | SGK_preRx_norı | SGK_preRx | SGK_Norm | EGFR L858R, KRAS-AMP | Pre Treatment | Pre_treatment | Pre_treatment | Primary tumor | N/A | | EGFR | EGFRp.L858R | EGFR | 1 | 130 | 81 | 49 | 123 | 4 | 3 | 0 |
| CB2 | Patient_6 | CB2_BX-1_norm | CB2_BX-1 | CB2_Norm | EGFR L861Q, EGFR L62R, EGFR R521K, EGFR UTR3 *16G>A, *32>A | Pre Treatment | Pre_treatment | Pre_treatment | Primary tumor | N/A | | EGFR | EGFRp.L62R, p.L861Q | EGFR | 0 | 56 | 40 | 16 | 52 | 1 | 2 | 1 |
| TH29 | Patient_7 | TH29_Pre_norm | TH29_Pre | TH29_Norm | EGFR L858R | Pre Treatment | Pre_treatment | Pre_treatment | Primary tumor | N/A | | EGFR | EGFRp.L747_P753delinsS | EGFR | 0 | 61 | 39 | 22 | 58 | 0 | 3 | 0 |
| TH210 | Patient_8 | TH210_E1_norn | TH210_E1 | TH210_Norm | ALK | treatment naive | Pre_treatment | Pre_treatment | abd mass | 0, none | Never | ALK | none | ALK | 0 | 127 | 65 | 62 | 113 | 0 | 12 | 2 |
| TH310 | Patient_9 | TH310E3_WB | TH310E3 | TH310WB | EGFR Exon 19 del | Pre Treatment | Pre_treatment | Pre_treatment | lobectomy | Surgical Resection | Never | EGFR | none | EGFR | 1 | 126 | 78 | 48 | 120 | 3 | 3 | 0 |
| TH231 | Patient_10 | TH231_E1_norn | TH231_E1 | TH231_Norm | ALK-EML4 | treatment naïve | Pre_treatment | Pre_treatment | LN | 0, none | Never | ALK | none | ALK | 0 | 47 | 21 | 26 | 46 | 0 | 1 | 0 |
| TH239 | Patient_11 | TH239E5_WB | TH239E5 | TH239WB | EGFR G719S E709A | Pre Treatment | Pre_treatment | Pre_treatment | pleural fluid | No Tx | Former | EGFR | none | EGFR | 0 | 70 | 16 | 54 | 15 | 26 | 29 | 0 |
| TH52 | Patient_12 | TH52_A1_pre_n | TH52_A1_p | TH52_Norm | EGFR L858R | Pre Treatment | Pre_treatment | Pre_treatment | Primary tumor | N/A | | EGFR | EGFRp.L858R | EGFR | 0 | 72 | 50 | 22 | 61 | 4 | 7 | 0 |
| TH266 | Patient_13 | TH-266-E2_norn | TH-266-E2 | TH266_Norm | ALK Fusion | Pre_treatment | Pre_treatment | Pre_treatment | | Chemo | | ALK | none | none | 0 | 25 | 14 | 11 | 22 | 2 | 1 | 0 |
| TH91 | Patient_14 | TH91_a_norm | TH91_a | TH91_Norm | EGFR L858R | Pre Treatment | Pre_treatment | Pre_treatment | Primary tumor | N/A | | EGFR | EGFRp.L858R, p.R108G | EGFR | 0 | 68 | 42 | 26 | 61 | 3 | 4 | 0 |
| TH176 | Patient_15 | TH176E2_WB | TH176E2 | TH176WB | EGFR Exon 19 del | Pre Treatment | Pre_treatment | Pre_treatment | right 12th rib | No Tx | Former | EGFR | none | EGFR | 0 | 218 | 145 | 73 | 213 | 0 | 3 | 2 |
| TH169 | Patient_16 | TH169_E2_norn | TH169_E2 | TH169_Norm | EGFR Exon 19 del | treatment naive | Pre_treatment | Pre_treatment | lung | None | Never | EGFR | none | EGFR | 0 | 93 | 51 | 42 | 84 | 1 | 7 | 1 |
| TH205 | Patient_17 | TH205_E2_norn | TH205_E2 | TH205_Norm | EGFR L858R | treatment naïve | Pre_treatment | Pre_treatment | pleura | none | Never | EGFR | none | EGFR | 1 | 69 | 39 | 30 | 66 | 1 | 2 | 0 |
| TH17 | Patient_18 | TH17_pre_norm | TH17_pre | TH17_Norm | EGFR Exon 19 del | Pre Treatment | Pre_treatment | Pre_treatment | Pleural Fluid | N/A | | EGFR | EGFRp.E746Ifs*16, p.R748Qfs*16 | EGFR | 0 | 43 | 35 | 8 | 40 | 0 | 3 | 0 |
| CB1 | Patient_19 | CB1_BX-1_norn | CB1_BX-1 | CB1_Norm | EGFR L858R, EGFR L62R | Pre Treatment | Pre_treatment | Pre_treatment | Primary tumor | N/A | | EGFR | EGFRp.L858R, p.L62R | EGFR | 1 | 118 | 75 | 43 | 103 | 8 | 7 | 0 |
| TH256 | Patient_20 | TH256E4_WB | TH256E4 | TH256WB | EGFR L858R | Pre Treatment | Pre_treatment | Pre_treatment | pleural fluid, cell button | No Tx | Never | EGFR | EGFRp.L62R | EGFR | 0 | 45 | 34 | 11 | 45 | 0 | 0 | 0 |
| TH203 | Patient_21 | TH203E4_WB | TH203E4 | TH203WB | EGFR Exon 19 del | Pre Treatment | Pre_treatment | Pre_treatment | Right upper lobe | No Tx | Never | EGFR | none | EGFR | 1 | 136 | 90 | 46 | 124 | 5 | 7 | 0 |
| TH148 | Patient_22 | TH148E1_WB | TH148E1 | TH148WB | EGFR Exon 19 del | Pre Treatment | Pre_treatment | Pre_treatment | Lung tumor | No Tx | Never | EGFR | none | EGFR | 1 | 114 | 73 | 41 | 109 | 2 | 2 | 1 |
| TH275 | Patient_23 | TH275E1_WB | TH275E1 | TH275WB | EGFR Exon 19 del | Pre Treatment | Pre_treatment | Pre_treatment | left side pleural fluid | No Tx | Never | EGFR | EGFRp.N158= | EGFR | 0 | 84 | 56 | 28 | 82 | 1 | 1 | 0 |
| TH087 | Patient_24 | TH087E5_WB | TH087E5 | TH087WB | EGFR Exon 19 del | Post Treatment 2 | Post_treatment | Multiple_therapies_TKI_last | Right adrenal mass | Osimertinib | Former | EGFR | EGFRp.E746_A750del | EGFR | 1 | 574 | 372 | 202 | 559 | 4 | 9 | 2 |
| AZ003 | Patient_1 | AZ03SURG_WE | AZ03SURG | AZ03WB | EGFR L858R | Post Treatment | Post_treatment | TKI | | Neoadj_osi | | EGFR | EGFRp.L858R | EGFR | 1 | 112 | 67 | 45 | 109 | 1 | 2 | 0 |
| TH239 | Patient_11 | TH239E4_WB | TH239E4 | TH239WB | EGFR G719S E709A | Post Treatment | Post_treatment_ | TKI | left LN2 | Erlotinib | | EGFR | EGFRp.E709A, p.G719S | EGFR | 0 | 244 | 93 | 151 | 122 | 63 | 59 | 0 |
| TH51 | Patient_25 | TH51_A2_norm | TH51_A2 | TH51_Norm | EGFR L858R | Post Treatment | Post_treatment | TKI | Liver Met | Erlotinib X 6 months PR | | EGFR | EGFRp.L858R | EGFR | 0 | 205 | 128 | 77 | 200 | 1 | 3 | 1 |
| TH067 | Patient_2 | TH67_E7_norm1 | TH067_E7 | TH067_Norm_1 | EGFR Exon 19 del | early rebiopsy (18 days on treatment) | Post_early | Multiple_therapies_TKI_last | liver (autopsy) | 6- chemo/ TKI/IO | Never | EGFR | EGFRp.E746Vfs*16, p.L747Ffs*14 | EGFR | 0 | 430 | 259 | 171 | 412 | 5 | 10 | 3 |
| TH171 | Patient_26 | TH171_E3_norn | TH171_E3 | TH171_Norm | EML4-ALK fusion Variant 3a/b | progressing disease | Post_treatment_ | Multiple_therapies_TKI_last | LN | 5- chemo/ tki/io | Never | ALK | ALKp.I1322M, p.E1210K | ALK | 0 | 220 | 128 | 92 | 216 | 0 | 4 | 0 |
| TH29 | Patient_7 | TH29_Post_norn | TH29_Post | TH29_Norm | EGFR L858R | Post Treatment 1 | Post_treatment | TKI | Liver Mass | Erlotnib X 6 months PD | | EGFR | none | EGFR | 0 | 14 | 7 | 7 | 12 | 1 | 1 | 0 |
| TH74 | Patient_27 | TH74_2_norm | TH74_2 | TH74_Norm | EGFR R521K | Post Treatment | Post_treatment | TKI | Lung Met | Erlotinib x 11 months PR | | EGFR | EGFRp.T790M, p.L747_P753delinsS | EGFR | 1 | 75 | 46 | 29 | 64 | 6 | 4 | 1 |
| AZ005 | Patient_28 | AZ05SURG_WE | AZ05SURG | AZ05WB | EGFR L858R | Post Treatment | Post_treatment_ | TKI | | Neoadj_osi | | EGFR | EGFRp.L858R | EGFR | 0 | 97 | 60 | 37 | 97 | 0 | 0 | 0 |
| AZ010 | Patient_29 | AZ10SURG_WE | AZ10SURG | AZ10WB | EGFR exon 19 del | Post Treatment | Post_treatment_ | TKI | | Neoadj_osi | | EGFR | none | none | 0 | 79 | 52 | 27 | 74 | 1 | 4 | 0 |
| TH330 | Patient_4 | TH330E1_WB | TH330E1 | TH330WB | EGFR L858R | Post Treatment | Post_treatment | TKI | No path | Osimertinib | Never | EGFR | none | EGFR | 1 | 148 | 98 | 50 | 141 | 0 | 7 | 0 |
| TH226 | Patient_30 | TH226_E3_norn | TH226_E3 | TH226_Norm | EGFR Exon 19 del | early rebiopsy (16 days) | Post_early | TKI | lung | | Never | EGFR | none | EGFR | 1 | 141 | 78 | 63 | 131 | 0 | 9 | 1 |
| TH153 | Patient_31 | TH153E3_WB | TH153E3 | TH153WB | EGFR Exon 19 del | Post Treatment | Post_treatment | TKI | retroperitoneal soft tissue | Erlotinib | Never | EGFR | EGFRp.T790M | EGFR | 0 | 124 | 80 | 44 | 111 | 2 | 10 | 1 |
| TH11 | Patient_32 | TH11_post_2_nc | TH11_post | TH11_Norm | EGFR Exon 19 del | Post Treatment 1 | Post_treatment | TKI | Pleural Fluid | Erlotinib X 3 months | | EGFR | EGFRp.E746_A750del, p.T790M | EGFR | 1 | 81 | 49 | 32 | 64 | 5 | 12 | 0 |
| TH155 | Patient_33 | TH155_E5_norn | TH155_E5 | TH155_Norm | EGFR Exon 19 del | progressing disease | Post_treatment_ | TKI | brain | 1, TKI | Never | EGFR | EGFRp.V1109Ifs*6 | EGFR | 1 | 84 | 40 | 44 | 77 | 1 | 6 | 0 |
| TH220 | Patient_34 | TH220_E2_norn | TH220_E2 | TH220_Norm | ALK-EML4 | progressing disease | Post_treatment | Multiple_therapies_TKI_last | liver | 5, TKI Chemo | Never | ALK | none | ALK | 0 | 142 | 93 | 49 | 133 | 1 | 7 | 1 |
| TH194 | Patient_35 | TH194E2_WB | TH194E2 | TH194WB | EGFR Exon 19 del, T790M | Post Treatment | Post_treatment | Multiple_therapies_TKI_last | pleural fluid | Osimertinib | Never | EGFR | none | EGFR | 0 | 16 | 8 | 8 | 10 | 1 | 5 | 0 |
| TH116 | Patient_36 | TH116_E2_norn | TH116_E2 | TH116_Norm | EGFR L858R | progressing disease | Post_treatment_ | TKI | liver | 2, TKI | Never | EGFR | EGFRp.L858R | EGFR | 1 | 125 | 68 | 57 | 122 | 1 | 2 | 0 |
| AZ009 | Patient_37 | AZ09SURG_WE | AZ09SURG | AZ09WB | EGFR exon 19 del | Post Treatment | Post_treatment | TKI | | Neoadj_osi | | EGFR | none | none | 1 | 97 | 62 | 35 | 93 | 1 | 2 | 1 |
| TH52 | Patient_12 | TH52_C_post_n | TH52_C_po | TH52_Norm | EGFR L858R | Post Treatment | Post_treatment | TKI | Lung Mass | Erlotinib X 8 years PR | | EGFR | EGFRp.L858R, p.T790M | EGFR | 0 | 124 | 93 | 31 | 113 | 1 | 10 | 0 |
| TH41 | Patient_38 | TH41_E2_norm | TH041_E2 | TH041_Norm | EML4-ALK V3a/b | progressing disease | Post_treatment | TKI | ovary | Chemo/TKI | Former | ALK | none | ALK | 0 | 109 | 62 | 47 | 107 | 0 | 2 | 0 |
| TH208 | Patient_39 | TH208_E4_norn | TH208_E4 | TH208_Norm | EGFR L861Q | early rebiopsy (24 days on tki) | Post_early | TKI | pleural fluid | none | Never | EGFR | EGFRp.L861Q | EGFR | 1 | 153 | 89 | 64 | 144 | 3 | 6 | 0 |
| TH169 | Patient_16 | TH169_E4_norn | TH169_E4 | TH169_Norm | EGFR Exon 19 del | on TKI 2.5 months | Post_treatment | TKI | lung | 1, TKI | Never | EGFR | none | EGFR | 0 | 116 | 65 | 51 | 106 | 3 | 6 | 1 |
| TH208 | Patient_39 | TH208_E3_norn | TH208_E3 | TH208_Norm | EGFR L861Q | progressing disease | Post_treatment_ | TKI | lung/ pleural fluid | | Never | EGFR | EGFRp.L861Q | EGFR | 1 | 130 | 82 | 48 | 120 | 2 | 8 | 0 |
| TH79 | Patient_3 | TH079E2_WB | TH079E2 | TH079WB | EGFR Exon 19 del; EGFR T790M | Post Treatment | Post_treatment | TKI | Right superclav metastatic carcinoma | Erlotinib | Never | EGFR | EGFRp.E746_A750del, p.T790M | EGFR | 1 | 320 | 207 | 113 | 300 | 11 | 9 | 0 |
| CB2 | Patient_6 | CB2_BX-2_norn | CB2_BX-2 | CB2_Norm | EGFR L861Q, EGFR L62R, EGFR R521K, EGFR UTR3 *16G>A, *32>A | Post Treatment | Post_treatment | TKI | | on first-line TKI | | EGFR | EGFRp.L861Q, p.L62R | EGFR | 0 | 37 | 30 | 7 | 31 | 2 | 4 | 0 |
| TH122 | Patient_40 | TH122E3_WB | TH122E3 | TH122WB | EGFR T790M, del19 | Post Treatment | Post_treatment_ | TKI | Lymph node, metastatic adenocarcinoma | Erlotinib | Never | EGFR | EGFRp.E746_A750del, p.T790M | EGFR | 0 | 145 | 101 | 44 | 130 | 6 | 8 | 1 |

**Supplementary Table 3:** Selective putative resistance mutations observed in *EGFR* - and *ALK* -driven lung cancer patients.

| **Patient_Specimen_ID** | **Alias** | **Oncogenic Driver** | **Putative resistance mutation(s) (APOBEC signature mutations/EGFR T790M)** | **Putative mechanism of resistance** | **Additional Comments** |
|---|---|---|---|---|---|
| AZ03 pre- and post-TKI | Patient_1 | EGFR L858R | RBM10 Q595* | Loss of RBM10, a splicing factor, has been shown to diminish EGFR inhibitor sensitivity by reducing ratio of BCL-Xs(pro-apoptotic) to BCL-XL (anti-apoptotic)[47]. | Had APOBEC signature mutations in major tumor suppressors including RBM10, Tp53 R213Q) and FBXW7 (S678*) indicating that APOBEC activity was likely activated early during tumorigenesis in this patient and may also be the driver of intrinsic resistance. |
| TH087_E5 | Patient_24 | EGFR exon19del | PTEN S287* and PIK3CA E545K | Activation of AKT signaling pathway[44, 39] | Inactivation of PTEN, a negative regulator of AKT signaling and activation of PIK3CA a major player in the AKT pathway would result in activation of AKT pathway. |
| TH51_A2 | Patient_25 | EGFR L858R | PPP2R2A S279* | Could reactivate MAPK pathway by inactivation of a negative regulator of Erk signaling, PP2A[40] | |
| TH11_post | Patient_32 | EGFR exon19del | EGFR T790M | Increased ATP affinity thereby diminishing the effect of EGFR inhibitor [37] | |
| TH122E3 | Patient_40 | EGFR exon19del | EGFR T790M | | |
| TH153E3 | Patient_31 | EGFR exon19del | EGFR T790M | | |
| TH52_C | Patient_12 | EGFR L858R | EGFR T790M | | |
| TH079E2 | Patient_3 | EGFR exon19del | EGFR T790M | | |
| TH171_E3 | Patient_26 | EML4-ALK | ALK 1210K | Reduces sensitivity to ALK inhibitors including Alectinib [41] | Also has ALK I322M mutation. |
| TH67_E7 | Patient_2 | EGFR exon19del | PPP2CA G247A, D160N, R135T, S93L and S43F | Could reactivate MAPK pathway by inactivation of a negative regulator of Erk signaling, PP2A[40] | |
| TH29 | Patient_7 | EGFR L858R | SMARCE1 E312K | Could reactivate EGFR signaling | Predicted to be functionally-impacting. SMARCE1 loss has been shown to diminish response to MET and ALK inhibition by induction of EGFR expression[46]. |
| TH239 | Patient_11 | EGFR E709A and G719S | PIK3CA E545K | Activation of AKT signaling pathway[44, 39] | |
